# Age-related differences in the neural network interactions underlying the predictability gain

**DOI:** 10.1101/2021.12.02.470763

**Authors:** Anna Uta Rysop, Lea-Maria Schmitt, Jonas Obleser, Gesa Hartwigsen

## Abstract

Speech comprehension is often challenged by increased background noise, but can be facilitated via the semantic context of a sentence. This predictability gain relies on an interplay of language-specific semantic and domain-general brain regions. However, age-related differences in the interactions within and between semantic and domain-general networks remain poorly understood. Using functional neuroimaging, we investigated commonalities and differences in network interactions enabling processing of degraded speech in healthy young and old participants. Participants performed a sentence repetition task while listening to sentences with high and low predictable endings and varying intelligibility. Stimulus intelligibility was adjusted to individual hearing abilities. Older adults showed an undiminished behavioural predictability gain. Likewise, both groups recruited a similar set of semantic and cingulo-opercular brain regions. However, we observed age-related differences in effective connectivity for high predictable speech of increasing intelligibility. Young adults exhibited stronger connectivity between regions of the cingulo-opercular network and between left insula and the posterior middle temporal gyrus. Moreover, these interactions were excitatory in young adults but inhibitory in old adults. Finally, the degree of the inhibitory influence between cingulo-opercular regions was predictive of the behavioural sensitivity towards changes in intelligibility for high predictable sentences in older adults only. Our results demonstrate that the predictability gain is relatively preserved in older adults when stimulus intelligibility is individually adjusted. While young and old participants recruit similar brain regions, differences manifest in underlying network interactions. Together, these results suggest that ageing affects the network configuration rather than regional activity during successful speech comprehension under challenging listening conditions.

## Introduction

In everyday conversations, speech comprehension is often hampered by challenging listening conditions, such as environmental noise. To maintain successful communication, listeners exploit facilitating factors, such as semantic context. The semantic context of a sentence provides information that can be used to predict upcoming words (“predictability gain”). This predictability gain has its maximum impact on comprehension at intermediate levels of intelligibility, that is, when the acoustic speech stream is neither completely intelligible nor completely unintelligible (Hartwigsen, Golombek, & Obleser, 2015; Pichora-Fuller, Schneider, & Daneman, 1995).

With respect to the neural correlates underlying the predictability gain, several neuroimaging studies found consistent activation in left (and less consistently right) angular gyrus (AG), left posterior middle temporal gyrus (pMTG) and left inferior frontal gyrus (IFG; Adank, 2012; Golestani, Hervais-Adelman, Obleser, & Scott, 2013; Obleser & Kotz, 2010; Obleser, Wise, Dresner, & Scott, 2007; Rysop, Schmitt, Obleser, & Hartwigsen, 2021). These regions have been suggested to constitute key regions of semantic processing (Binder, Desai, Graves, & Conant, 2009; Jefferies, 2013; Seghier, 2013). In addition, some studies also showed increased connectivity of left angular gyrus with other nodes of the semantic network when listening to high versus low predictable sentences at intermediate levels of intelligibility (Golestani et al., 2013; Obleser & Kotz, 2010; Obleser et al., 2007). Other work demonstrated the functional relevance of left angular gyrus for the predictability gain with inhibitory neurostimulation (Hartwigsen et al., 2015). Together, these studies provide evidence for a key role of semantic regions, particularly left angular gyrus, in the successful integration of semantic information into sentential context under challenging listening conditions in healthy young adults.

Aside from the contribution of semantic regions, sentence comprehension under challenging listening conditions results in increased executive demands (Fitzhugh, Schaefer, Baxter, & Rogalsky, 2021), leading to a recruitment of domain-general networks. Specifically, the cingulo-opercular network (Dosenbach, Fair, Cohen, Schlaggar, & Petersen, 2008; Uddin, Yeo, & Spreng, 2019), encompassing bilateral anterior insulae as well as a portion of the dorsal anterior cingulate and pre-supplementary motor area (pre-SMA), is the most frequently reported network in degraded speech processing (Alain, Du, Bernstein, Barten, & Banai, 2018; Erb & Obleser, 2013; Rogers & Peelle, 2022; Vaden Jr., Kuchinsky, Ahlstrom, Dubno, & Eckert, 2015; Vaden Jr. et al., 2013). Prior studies investigated the contribution of cingulo-opercular regions to challenging listening conditions either by using manipulations of the auditory signal itself (e.g., noise vocoding, where spectral information of an acoustic signal is degraded while temporal information is preserved, see Shannon, Zeng, Kamath, Wygonski, & Ekelid, 1995) or by masking the acoustic signal with broadband noise or multi-talker babble (Darwin, 2008; Miller, 1947). Although increased activity in cingulo-opercular regions has been linked to error monitoring, executive functions and maintenance of auditory attention (Wilsch, Henry, Herrmann, Maess, & Obleser, 2015), its exact role in acoustically-demanding speech perception remains vaguely defined. In one study, increased activity in cingulo-opercular regions during challenging listening conditions was predictive of successful word recognition in the following trial (Vaden et al. 2013), thereby indicating that activity in these regions is directly relevant to behaviour. In line with this observation, Rysop et al. (2021) investigated context-dependent changes in effective connectivity between nodes of the cingulo-opercular network during adverse listening conditions. They found a stronger inhibitory influence on the connections within the cingulo-opercular network, when semantic context was highly predictive and intelligibility increased. Crucially, the degree of inhibitory influence at intermediate levels of intelligibility was associated with an increased behavioural predictability gain. This finding suggests that inhibition of the cingulo-opercular network is beneficial in young adults, as long as semantic cues can be efficiently used to support comprehension of the distorted signal. Otherwise, increased activity in cingulo-opercular regions might help to maintain task performance. Yet, the contribution of the cingulo-opercular network to challenging listening conditions in the ageing brain is unclear.

Ageing is associated with a decline across sensory and cognitive domains such as hearing acuity, processing speed or working memory capacity (Salthouse, Atkinson, & Berish, 2003; Wingfield, Amichetti, & Lash, 2015). However, semantic abilities are thought to remain relatively stable across the lifespan (Verhaeghen, 2003; Wingfield & Stine-Morrow, 2000). While listening to degraded speech is already demanding for young listeners, older adults frequently report enhanced difficulties in understanding speech in challenging listening situations, even if they show no signs of age-related hearing loss or difficulties in ideal listening conditions (Gordon-Salant, 2005; Gordon-Salant & Fitzgibbons, 1995; Humes, 1996; Pichora-Fuller, Alain, & Schneider, 2017). With regard to the predictability gain, behavioural studies point towards either an increased reliance (Desjardins & Doherty, 2013; Goy, Pelletier, Coletta, & Kathleen Pichora-Fuller, 2013; Pichora-Fuller et al., 1995; Sheldon, Pichora-Fuller, & Schneider, 2008) or similar use of contextual information in older compared to younger adults (Dubno, Ahlstrom, & Horwitz, 2000; Sheldon et al., 2008). For example, in a semantic priming paradigm with an acoustically degraded prime, older participants benefitted from contextual information as much as young participants (Sheldon et al. 2008).

While the beneficial effect of semantic context generally seems to be preserved in older adulthood, changes at the neural level remain debated. Some authors argue that the reduced magnitude of electrophysiological measures to high predictable sentences in old versus young adults might reflect less efficient use of sentential context (Federmeier, 2007; Payne & Federmeier, 2018; Wlotko, Federmeier, & Kutas, 2012). Likewise, a recent meta-analysis showed less activation in semantic key regions for older participants but increased recruitment of domain-general regions during semantic tasks (Hoffman & Morcom, 2018). Yet, these changes were mainly observed when task performance was poor. Thus, it remains unclear whether increased recruitment of domain-general networks in the ageing brain reflects a (compensatory) attempt to maintain performance during speech comprehension under challenging listening conditions (Eckert et al., 2008; Peelle, Troiani, Grossman, & Wingfield, 2011) or rather a pattern of dedifferentiation, indicating less functional specialization (Li, Lindenberger, & Sikström, 2001). Some authors suggest that older adults already recruit such domain-general resources under relatively easy conditions (Erb & Obleser, 2013; Fitzhugh, Braden, Sabbagh, Rogalsky, & Baxter, 2019; Peelle, 2018; Reuter-Lorenz & Cappell, 2008). Accordingly, stronger activity in domain-general regions was associated with better comprehension of degraded auditory sentences selectively in old adults (Erb & Obleser, 2013). These findings suggest that the (potentially compensatory) upregulation of activity in domain-general areas is more pronounced in older listeners.

The additional upregulation of domain-general regions is accompanied by altered patterns of functional connectivity (Sala-Llonch, Bartres-Faz, & Junque, 2015). It has been consistently reported that older adults exhibit decreased levels of connectivity within domain-general networks including the cingulo-opercular network (Geerligs, Renken, Saliasi, Maurits, & Lorist, 2015), while connectivity between networks was increased at rest (Setton et al., 2022; Zonneveld et al., 2019) and during cognitive tasks (Zhang, Gertel, Cosgrove, & Diaz, 2021). However, studies addressing age-differences in task-related connectivity during language processing are sparse. Hence, it remains unclear how ageing affects functional interactions within and between networks during speech comprehension under adverse listening conditions.

In the present study, we assessed age-related differences in the behavioural predictability gain and its neural correlates and network configuration while participants listened to sentences of varying acoustic intelligibility and semantic predictability. In contrast to previous studies, we used an optimized adaptive tracking paradigm that considered individual differences in auditory perception by adjusting stimulus intelligibility to individual hearing abilities. This assures comparable task performance across participants, which is particularly important in light of variable hearing performance in old participants. Using dynamic causal modelling (DCM) we analysed age-related differences in connectivity within the semantic and cingulo-opercular networks, respectively (i.e. within-network connectivity) as well as between nodes of both networks (i.e. between-network connectivity).

We expected a beneficial effect of semantic context when speech is acoustically degraded in both age groups, as intelligibility levels were adjusted to individual hearing abilities. At the neural level, we expected potentially weaker engagement of semantic regions and relatively stronger recruitment of cingulo-opercular regions in older listeners. Age-related differences should also be reflected in decreased connectivity within the cingulo-opercular and semantic network and increased connectivity between the semantic and cingulo-opercular network in older listeners, reflecting increased recruitment of domain-general resources.

To foreshadow our results, when controlling for speech-to-noise reception thresholds as done in the present study, no age-related differences in the behavioural performance or task-related activity were observed. However, relative to young listeners, older listeners showed decreased effective connectivity within the cingulo-opercular network and between left anterior insula and posterior middle temporal gyrus when predictability was high and intelligibility increased. Stronger inhibitory influence between cingulo-opercular regions was associated with better task performance selectively in older listeners, demonstrating its behavioural relevance.

## Materials and Methods

### Participants

Thirty healthy middle-aged to old German native speakers were recruited for the fMRI experiment. The sample size was determined based on our previous study (Rysop et al., 2021). Inclusion criteria comprised age-normal hearing (tested with pure-tone audiometry; see below for further details) and right-handedness according to the German version of the Edinburgh Handedness Inventory (Oldfield, 1971). Participants were excluded if they showed signs of cognitive impairment (Mini Mental State Examination score < 27; MMSE; Folstein, Folstein, & McHugh, 1975) or reported a history of neurological or psychiatric disorders. Data from three participants were excluded from the analyses due to excessive head movement during fMRI (movement parameter exceeded 1.5 times the voxel size), and data from one participant because of a ceiling effect in the behavioural performance.

The final sample size was *n* = 26 (M = 62 years, range: 50–77 years, 19 female, see Supplementary Table 1 for further information). This group was compared to 26 healthy young participants (M = 25 years, range: 19–29 years, 15 female), who had performed the same in-scanner task under the exact same procedures in an earlier experiment (see Rysop et al., 2021 for a detailed analysis of the neural predictability gain in this younger group). Participants were reimbursed with 10 € per hour and gave written informed consent in accordance to the declaration of Helsinki prior to participation. The study was approved by the local ethics committee (University of Leipzig).

### Experimental Procedure

The experiment consisted of two sessions. In the first session, each participant’s hearing status and cognitive abilities were tested using pure-tone audiometry and MMSE. Only participants with relatively well-preserved peripheral hearing abilities (pure-tone average < 25 dB HL in the listener’s better ear) were included. The hearing test was conducted in a soundproof chamber using an audiometer (Oscilla SM910-B Screening Audiometer). Pure-tone averages were computed across frequencies from 250 kHz to 8000 kHz in the right and left ear separately (see Figure 1A for an illustration of the peripheral hearing status of included participants). Six participants had a PTA > 25 dB in one ear, but met the criterion of PTA < 25 dB in the other ear and were therefore included. For included participants, the first session proceeded with further neuropsychological assessments. Working memory was tested using the auditory forward and backward version of the Digit Span Task (Wechsler, 1955). Measures of executive functions (processing speed and cognitive flexibility) were assessed with both versions of the Trail Making Test (Reitan, 1958). Further, we tested verbal fluency and flexibility using two subtests of a German word fluency test (Regensburger Wortflüssigkeitstest; Aschenbrenner, Tucha, & Lange, 2000). In this test, participants had to generate as many nouns as possible either for one specific semantic category (subtest A, category *food*) or two alternating semantic categories (subtest B, categories *fruits* and *sports*) within two minutes. The first session had a duration of approximately one hour.

**Figure 1.**
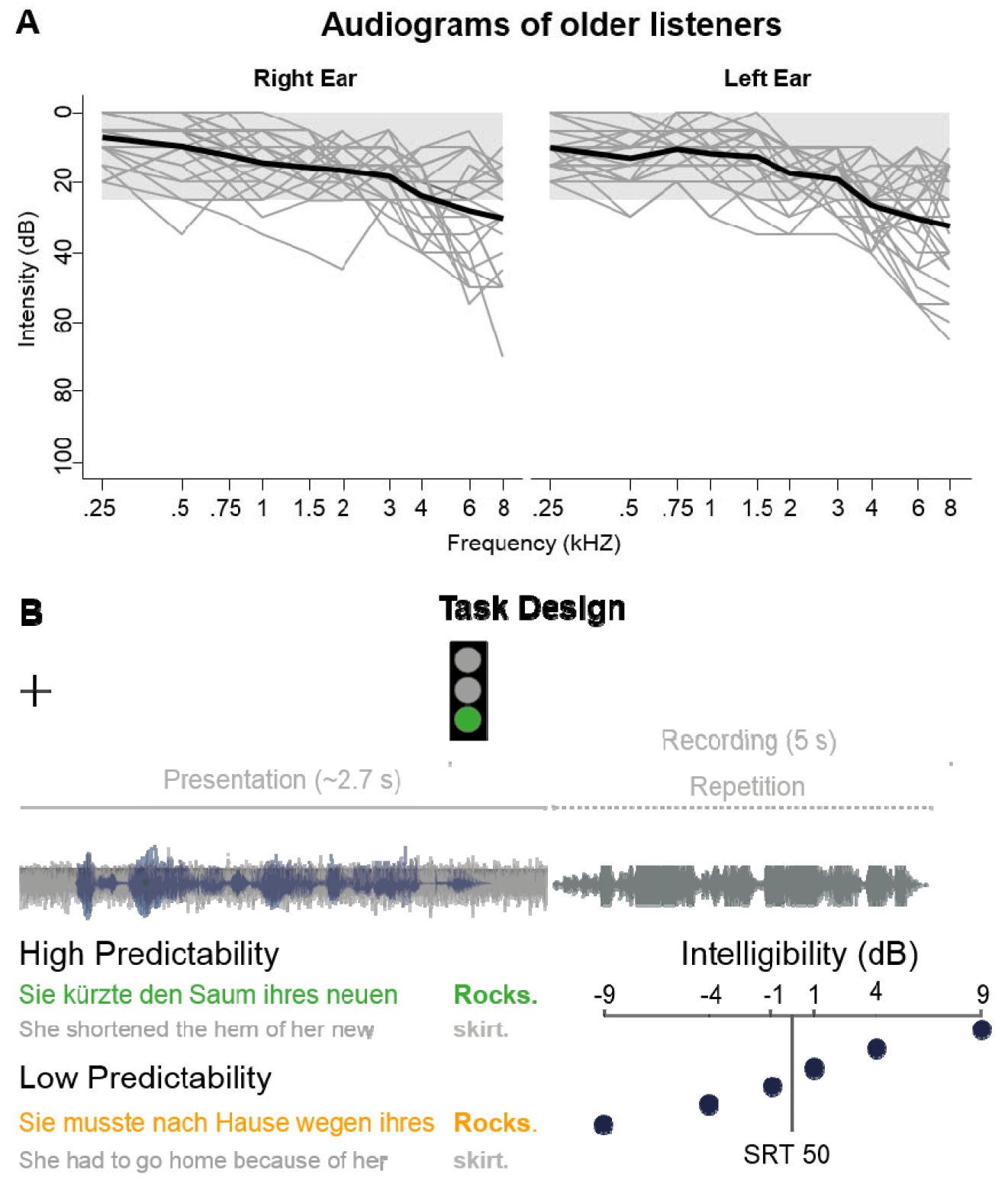
Experimental procedures. **A**. Pure-tone audiograms for the older participants, plotted for the right and left ear separately. Grey lines represent the hearing curve of single participants across each tested frequency; the bold black line represents the average across participants. The intensity range for normal hearing is shaded in grey. **B**. In the sentence repetition task, each trial consisted of a listening period in which a short sentence (blue waveform) was embedded in speech-shaped noise (grey waveform) and a fixation cross was presented on the screen. With the onset of the sentence-final word (i.e., keyword), a green traffic light appeared and marked the recording period, in which participants were asked to repeat the whole sentence as accurately as possible. The sentence context was either highly (green) or lowly (orange) predictive of the sentence-final keyword. The signal-to-noise ratio of the sentences against the speech-shaped background noise was adjusted to individual speech reception thresholds (SRT 50).

The fMRI experiment was conducted in a second session on a separate day. Participants were placed in the scanner and equipped with MR compatible in-ear headphones (MR Confon; Magdeburg, Germany). The volume was initially set to a comfortable intensity. To determine the participants’ in-scanner speech reception threshold (SRT; i.e., the signal-to-noise ratio (SNR) required to correctly repeat a sentence with a probability of 50 %), we conducted an adaptive up-down staircase experiment (Kollmeier, Gilkey, & Sieben, 1988; for a detailed description of the procedure see Rysop et al., 2021). The SRT was used as a reference to adjust stimulus intelligibility for each participant individually in the main experiment, thereby correcting for differences in peripheral hearing and ensuring comparable task performance across participants.

In the main experiment, participants listened to sentences while viewing a white fixation cross on a black screen. Participants were instructed to overtly repeat the whole sentence as accurately as possible to prevent listening strategies such as focussing on the last word. If they had not understood anything, they were asked to say so. Responses were cued by a green traffic light that appeared on the screen (see Figure 1B for an overview of the experimental design). Overt responses were recorded for a period of 5 seconds using a FOMRI-III microphone (Optoacoustics, Yehuda, Israel). The intertrial interval varied between 2000 and 7000 ms and served as an implicit baseline for the fMRI analysis. Stimuli were presented in six blocks using Presentation (version 18.0, Neurobehavioral Systems, Berkeley, USA, www.neurobs.com). The fMRI experiment had a duration of 50 minutes, the total duration of the second session was approximately 90 minutes.

### Stimulus Material

Experimental sentences were taken from the German Speech Intelligibility in Noise (G-SPIN) material (Kalikow, Stevens, & Elliott, 1977; for a detailed description of the German version, see Erb, Henry, Eisner, & Obleser, 2012). The 216 sentences consisted of pairs with the same sentence-final words (i.e., keyword) but different preceding sentence frames: While the frame of one of the sentences was predictive of the keyword (high predictability: *“She shortened the hem of her new <u>skirt</u>*.”), the other one was not (low predictability: *“She had to go home because of her <u>skirt</u>”*). Twenty additional G-SPIN sentences with highly predictable keywords were used for the adaptive tracking procedure.

The intelligibility of the sentences was manipulated by varying the sentence intensity relative to constant speech-shaped noise in the background. Sentence intensity was varied symmetrically and relative to the previously determined SRT in six steps (−9 dB, -4 dB, -1 dB, +1 dB, +4 dB, +9 dB SNR relative to the individual SRT; Rysop & Schmitt et al. 2021 for details). These intelligibility levels were chosen to cover the full range of behavioural performance in each participant while sampling those (intermediate) intelligibility levels with the largest predictability gain more densely.

Experimental conditions yielded a factorial design with the factors *predictability* (high, low) and *intelligibility* (−9 dB, -4 dB, -1 dB, +1 dB, +4 dB, +9 dB SNR relative to SRT), with 18 sentences per condition. Sentences were presented in a pseudorandomized order with the restriction that each intelligibility level was not presented more than three times in a row to prevent adaptation.

### MRI acquisition

Functional MRI data were collected on a 3 Tesla Siemens Prisma Scanner with a 32-channel head coil. We used a dual gradient-echo planar imaging multiband sequence (Feinberg et al., 2010). The scanning parameters were: TR = 2,000 ms; TE = 12 ms, 33 ms; flip angle = 90°; voxel size = 2.5 × 2.5 × 2.5 mm with an interslice gap of 0.25 mm; FOV = 204 mm; multiband acceleration factor = 2. For each participant, 1,500 volumes à 60 slices were acquired in axial direction and interleaved order. Slices were tilted by 10° off the AC-PC line to increase coverage of anterior temporal lobe regions. Field maps were acquired for later distortion correction (TR = 620 ms; TE = 4 ms, 6.46 ms). Additionally, high-resolution T1-weighted images were either obtained from the in-house database if available or were acquired in the second session using an MPRAGE sequence (whole brain coverage, TR = 1300 ms, TE = 2.98 ms, voxel size = 1 × 1 × 1 mm, matrix size = 256 × 240 mm, flip angle = 9°).

## Data Analysis

### Behavioural Analyses

All participants’ spoken response recordings were cleaned from scanner noise using Audacity (version 2.2.2, https://www.audacityteam.org/) and transcribed by one rater, who additionally determined speech onset times and speech durations. Repetitions of the keywords were rated either as correct or incorrect before calculating the proportion of correctly repeated keywords per participant and condition. Thereafter, psychometric curves were fitted to keyword repetition accuracies across intelligibility levels, separately for sentences with low and high predictability. More specifically, we employed cumulative Gaussian sigmoid functions using the Psignifit toolbox (Fründ, Haenel, & Wichmann, 2011) in MATLAB (version R2018b, MathWorks; see Supplementary Material for further details).

All empirical deviances between the fitted and a saturated model fell within the 97.5% confidence interval of their respective reference distribution, indicating that psychometric curves properly represented the observed behavioural data in every participant. There was no significant difference between the deviance of fits for sentences with low vs. high predictability (t_24_ = 0.11, p = .916, r = 0.06, BF10 = 0.21). To illustrate goodness of fit, we additionally calculated the R^2^_KL_ which is based on the information-theoretic measure of Kullback-Leibler divergence and represents the reduction of uncertainty by the fitted model relative to a constant model (Cameron & Windmeijer, 1997). On average, psychometric curves of sentences with high predictability yielded an R^2^_KL_ of 0.96 (SD = 0.04, range: 0.86– 0.995), sentences with low predictability an R^2^_KL_ of 0.94 (SD = 0.05, range: 0.81–0.996).

To quantify the predictability gain, we compared threshold and slope parameters of the psychometric functions for sentences with low versus high predictability. The threshold parameter represents the intelligibility level at which participants correctly repeat half of the keywords. A smaller threshold parameter for sentences with high compared to low predictability would indicate a stronger benefit from semantic context already at levels of lower intelligibility. The slope parameter describes the steepness of the curve at a proportion correct of 0.5 and denotes the sensitivity to changes in intelligibility at intermediate levels. Parameter estimates of the two psychometric curves were analysed using an ANOVA with the factors *age* and *predictability*. Furthermore, we analysed speech onset times (SOTs) as a measure of response speed. SOTs were log-transformed and submitted to a three-way ANOVA with the factors *age, predictability* and *intelligibility*. Statistical analyses were performed with RStudio (version 4.0.2; R Core Team, 2021) and JASP (version 0.9.1; https://jasp-stats.org/; Wagenmakers et al., 2018).

### Functional MRI analyses

Functional MRI data were preprocessed and analysed in SPM12 (version 7219, Wellcome Department of Imaging Neuroscience, London, UK) and MATLAB (version R2018b). As a first step, the images of the two echo times were combined based on a weighted average of the voxel-wise temporal signal-to-noise ratio maps of the first 30 volumes and realigned using a custom Matlab script. The rationale behind this approach is to improve signal quality in brain regions that typically suffer from signal loss (e.g., anterior temporal lobes; see Halai, Welbourne, Embleton, & Parkes, 2014 for a similar approach). Further, fMRI data were distortion corrected, co-registered to individual high-resolution structural T1-scans and spatially normalized to the standard template by the Montreal Neurological Institute (MNI). We preserved the original voxel size and finally smoothed the data, using a 5 mm^3^ FWHM Gaussian kernel to allow statistical inference based on Gaussian random-field theory (Friston, Worsley, Frackowiak, Mazziotta, & Evans, 1994).

Preprocessed data were submitted to a general linear model (GLM). Sentence presentation times were modelled with stick functions and convolved with SPM’s canonical hemodynamic response function. We included one regressor of no interest capturing the verbal responses using participant- and trial-specific speech onset times and durations. As additional nuisance regressors we included the six motion parameters obtained from the rigid-body transformation of the realignment step (see Supplementary Figure 1) and a vector for each volume that exceeded a framewise displacement of 0.9 mm (Siegel et al., 2014). A high-pass filter with a cut-off at 128 s was applied to the data.

At the single-participant level of both age groups, we computed direct contrasts between experimental conditions. These contrasts encompassed the main effect of intelligibility (i.e., linear increase in intelligibility, –9 < –4 < –1 < +1 < +4 < +9 dB SNR), the main effect of predictability (high predictable > low predictable sentences and vice versa) and the interaction between predictability and intelligibility (high predictable > low predictable sentences, with a linear increase in intelligibility and vice versa). The single-participant interaction contrasts were further used to extract the timeseries for the effective connectivity analysis.

The resulting contrast maps were submitted to one-sample t-tests, to investigate effects **within** each age group. Two-sample t-tests were used to investigate differences **between** groups (Poldrack, Nichols, & Mumford, 2011). To characterize commonly activated regions by both groups, the respective contrast maps were submitted to a conjunction analysis based on the minimum statistic (Nichols, Brett, Andersson, Wager, & Poline, 2005). To correct for multiple comparisons, we used the family-wise error (FWE) technique applying a cluster-level threshold of *p* < 0.05 and a voxel-wise threshold of *p* < 0.001). Statistical maps were visualized with BrainNet Viewer (Xia, Wang, & He, 2013). Anatomical locations were specified using the SPM anatomy toolbox (version 3.0; Eickhoff et al., 2005, 2007) and the Harvard-Oxford atlas (https://fsl.fmrib.ox.ac.uk/fsl/fslwiki/).

### Dynamic Causal Modelling (DCM)

We used Dynamic Causal Modelling (DCM; version 12.5; Friston, Harrison, & Penny, 2003) and Parametric Empirical Bayes (PEB; Zeidman, Jafarian, Corbin, et al., 2019; Zeidman, Jafarian, Seghier, et al., 2019) to investigate age-related differences in effective connectivity. DCM is a method that uses a biophysically informed generative model to estimate effective connectivity between a set of brain regions. The estimated signal is compared to the measured fMRI BOLD signal and quantified in terms of negative variational free energy. Three sets of parameters have to be defined: (1) intrinsic parameters, reflecting endogenous connectivity between regions in the absence of experimental manipulation (A-Parameters), (2) modulatory parameters that encode the rates of change in connectivity due to an experimental manipulation (B-Parameters) and (3) driving input parameters, reflecting the influence of an experimental task on single nodes within the network (C-Parameters).

While the classical DCM approach allows to investigate the network architecture (or configuration) underlying an experimental effect, PEB spans a hierarchical model over the connectivity parameters, allowing to quantify differences and commonalities in coupling strengths among groups of participants (Friston et al., 2016; Zeidman, Jafarian, Corbin, et al., 2019). PEB provides a Bayesian GLM-like approach that divides inter-subject variability into regressor effects and unexplained random effects (Zeidman, Jafarian, Seghier, et al., 2019). In contrast to classical inference over parameter estimates, the PEB framework does not only take the mean but also the uncertainty of individual parameters into account. Consequently, participants with more uncertain parameters will influence the group estimates less than participants with more certain parameters (Zeidman, Jafarian, Seghier, et al., 2019).

We were interested in the context-dependent effective connectivity *within* the semantic and cingulo-opercular networks as well as *between* nodes of both networks. We approached these questions in a two-step procedure. First, we performed a standard DCM analysis within the cingulo-opercular network, motivated by the findings of our previous study, to test the goodness of fit of the winning model that was identified in young participants (Rysop et al., 2021). In that study, the data of young participants was best explained by a model in which intelligibility and predictability of the sentences jointly modulated the connections from pre-SMA and right anterior insula to the left anterior insula. This model had a relatively high exceedance probability of 0.73. Here, we aimed at testing whether the optimal model for young participants also succeeds in explaining the older participants’ data (see Supplementary Methods for a detailed description of the standard DCM approach).

Second, we employed the DCM PEB approach which enabled us to test for age-related effects in a larger model space encompassing nodes of the semantic and cingulo-opercular network identified by the conjunction analysis. Specifically, we asked how predictability and intelligibility of sentences affects the coupling (1) within the semantic and cingulo-opercular network, (2) between both networks, as well as (3) whether coupling strengths differ across groups and (4) whether age-related differences in coupling strength are related to behavioural differences.

#### Seed Region Selection

For the standard DCM approach, we defined the seed regions of the cingulo-opercular network based on the interaction contrast in the older subgroup. The seed regions comprised the pre-SMA [-4 20 42] and anterior insulae [left: -30 23 -2, right: 33 26 -2]. For the PEB DCM approach, we defined the seed regions based on the results of the conjunction analyses and on our previous findings in young participants (Rysop et al., 2021). Seed regions comprised bilateral AG [left: -42 -70 35; right: 46 -64 35] and left pMTG [-57 -60 -2] of the semantic network, as well as pre-SMA [-7 13 48] and anterior insulae [left: -33 23 -2, right: 33 23 0] of the cingulo-opercular network. The first principal component (eigenvariate) was extracted from all voxels within a spherical region of interest (6 mm radius) that passed a liberal threshold of p_uncorrected_ < .05 for each seed region. The sphere was initially centred on the group coordinate and was allowed to move within a spherical search space of 10 mm around the group coordinate to account for variation in the exact participant-specific maximum. The search space for the bilateral insulae was constrained by a 5 mm sphere centred on the group coordinate to ensure that the individual peak fell within the anterior insula and not the surrounding regions. The extracted time courses were adjusted by an *F*-contrast spanning across the experimental conditions to regress out effects of no interest (i.e., overt speech or head movement).

#### PEB DCM Setup

The participant-level design matrix for the PEB DCM analysis consisted of three regressors. The first regressor coded the onsets of all experimental stimuli and served as driving input regressor. The ensuing regressors served as modulatory input and coded the effect of 1) high and 2) low predictability with the level of intelligibility added as parametric modulation (i.e., the effect of predictability, scaled linearly with intelligibility).

At the first-level, we specified and inverted one “full” DCM for each participant (see Figure 6A for a schematic full model). This model included all possible reciprocal connections between seed regions as well as self-connections. Because we did not include a primary sensory region in the model, we set the driving input to every seed region (C-Matrix). Each connection, except for the self-connections, could receive modulation by experimental inputs (B-Matrix). The inputs were not mean-centered, so that the intrinsic parameter estimates (A-matrix) can be interpreted as baseline connectivity in the absence of a task (Dijkstra, Zeidman, Ondobaka, Van Gerven, & Friston, 2017; Marreiros, Kiebel, & Friston, 2008). Note that parameter estimates represent rates of change of the influence from one region to another (intrinsic connectivity) or of an experimental manipulation on the connection between regions (modulatory connectivity). Positive parameters translate to a positive influence of one region onto another region or an experimental manipulation on a connection and can be interpreted as excitatory. Negative parameter estimates can be interpreted as inhibitory influence (Dijkstra et al., 2017; Zeidman, Jafarian, Corbin, et al., 2019; Zeidman, Jafarian, Seghier, et al., 2019). All full models were inverted using variational Laplace. We extracted the percent variance explained to obtain a parameter for the model fit.

At the second-level, we set up the PEB model of between-participant effects to investigate commonalities and differences in intrinsic and modulatory parameters. The PEB model included a group-level design matrix with one regressor encoding the mean across both groups (or commonalities across participants) whereas the second regressor encoded age group, to identify differences in connectivity between younger and older participants. In contrast to traditional DCM, this procedure does not compare the model evidence of different network architectures, but the effect of the presence or absence of an experimental manipulation on a connection. Thus, each connection, opposed to the whole network architecture, receives a posterior probability. To only keep those parameters with a high probability of contributing to the model evidence, we performed an automatic search over reduced models using Bayesian Model Comparison and Bayesian Model Reduction (BMR; Friston et al., 2016). In this procedure, parameters are systematically switched on and off (i.e., nested models) and compared against the full model, pruning away those parameters that do not contribute to the model evidence (negative variational free energy).

Lastly, we computed a Bayesian Model Average (BMA) over the parameters of the final iteration of the automatic search. Here, parameters were weighted by their posterior probabilities (Zeidman, Jafarian, Seghier, et al., 2019). To retain those parameters with ‘strong’ evidence of being non-zero (Kass & Raftery, 1995), we applied a threshold at a posterior probability of > 95 % based on free energy. Results were considered as within-network connectivity when task-dependent modulations were found between nodes of the semantic or the cingulo-opercular network, respectively. In case the task modulated the connection between one node of the semantic and one node of the cingulo-opercular network, this result was interpreted as between-network connectivity. Further, we probed whether the effect sizes were large enough to predict a left-out participant’s age group using leave-one-out cross-validation on the estimated parameters that had a significant age-related difference. Finally, to relate differences in connectivity to the behavioural performance, we extracted participant-specific parameters from the connections that showed a significant group effect and correlated these with the respective behavioural parameters of the psychometric curves.

## Results

### Behavioural Results

#### Semantic context benefits comprehension under challenging listening conditions independent of age

The average speech reception threshold obtained during the adaptive tracking procedure was significantly higher for older (3.75 dB) than for younger participants (1.68 dB; *t*(41.12) = 2.36, *p* = 0.023; Figure 2A). This finding is in line with a previous report that SRT’s are approximately 2 dB higher in older adults (Schneider, Pichora-Fuller, & Daneman, 2010).

**Figure 2.**
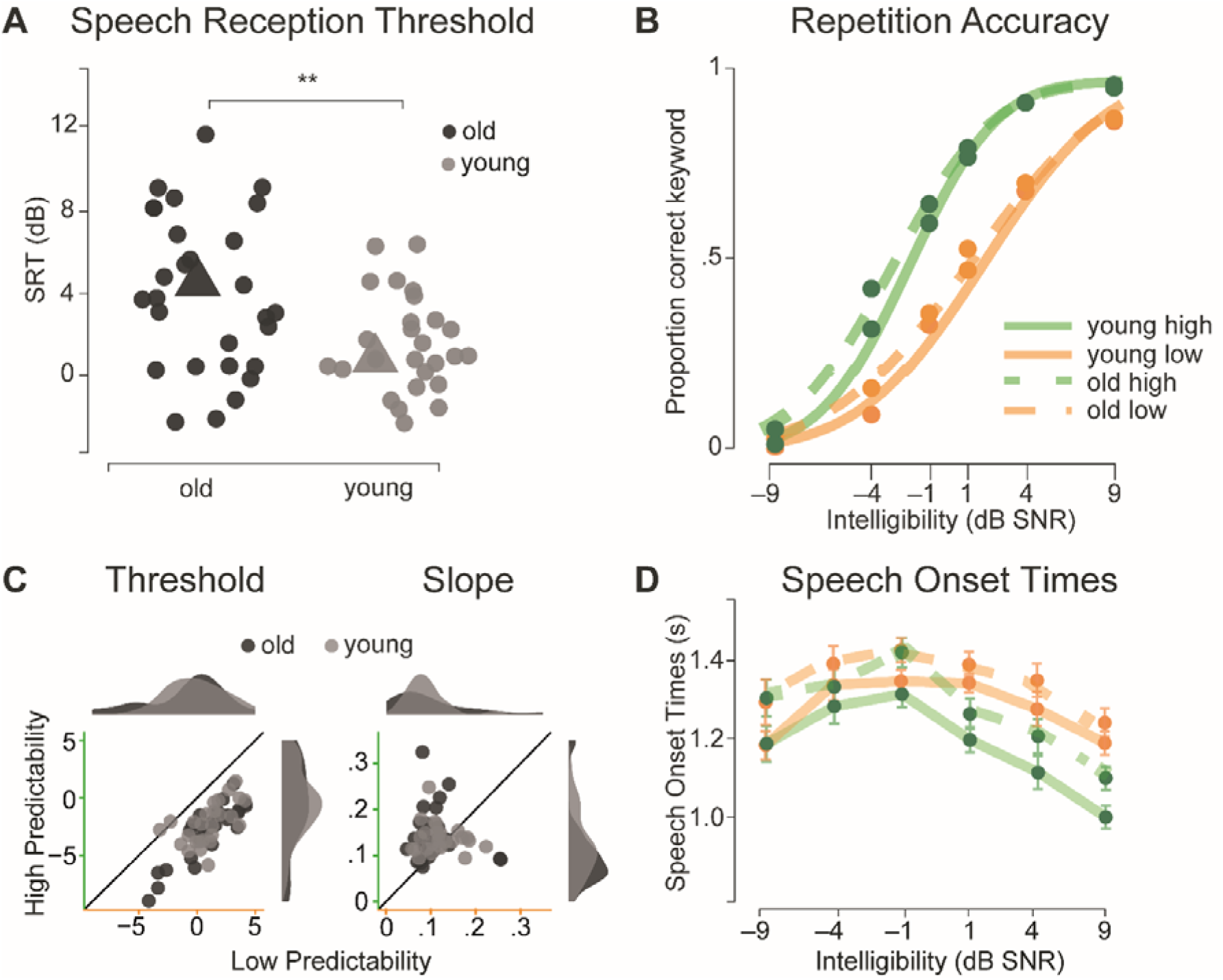
Behavioural results. **A** Speech reception thresholds (SRT) for old and young participants. Single dots represent individual participants; triangles represent the mean SRT per age group ** = *p* < 0.01. **B** Psychometric curves fitted to the proportion of correctly repeated keywords (dots) across intelligibility levels for sentences of high (green) and low (orange) predictability, separately for each age group (solid lines = young, dotted lines = old participants). **C** Individual threshold (left) and slope (right) parameters extracted from the psychometric curves at 50 % correct for high (y-axis) and lowly (x-axis) predictable keywords. Dark grey = old participants, light grey = young participants. **D**. Mean speech onset times for highly (green) and lowly (orange) predictable sentences across intelligibility levels.

On average, keyword repetition accuracies were comparable between young (M = 52.33 %, SD = 8.09 %) and old participants (M = 54.59 %, SD = 10.56 %). To explore effects of age on speech comprehension, we fitted psychometric curves to keyword repetition accuracies across intelligibility levels separately for both age groups and predictability levels (Figure 2B). We found a significant main effect of predictability for the threshold parameter of psychometric curves (*F*(1,50) = 302.3, *p* < 0.001, η^2^ = 0.41; Figure 2C), indicating that high predictability benefits comprehension already at lower levels of intelligibility. Further, a significant main effect of predictability on the slope parameters (*F*(1,50) = 7.46, *p* = 0.009, η^2^ = 0.08; Figure 2C) indicates that an increase in intelligibility benefits comprehension more strongly when speech is highly predictable. Together these results demonstrate that speech comprehension at intermediate levels of intelligibility generally benefits from high predictability. Most central to the aims of the present study, psychometric curves were strikingly similar between both age groups. Accordingly, the main effect of *age* and the interaction of *age* and *predictability* was not significant for any parameter of the psychometric curves (Supplementary Table 2 for all results).

The analysis of log-transformed speech onset times revealed a significant main effect of predictability (*F*(1,50) = 163.51, *p* < 0.001, η = 0.038), a significant main effect of intelligibility (*F*(5,250) = 36, *p* < 0.001, η = 0.08) and a significant interaction between predictability and intelligibility (*F*(5,250) = 23.29, *p* < 0.001, η = 0.02). Post-hoc pairwise comparisons showed that highly predictable sentences were repeated significantly faster than lowly predictable sentences at the three highest intelligibility levels (+1 dB SNR: *t*(101.54) = -2.997, *p*_corr_ = 0.003; +4 dB SNR: *t*(99.3) = -3.57, *p*_corr_ < 0.001; +9 dB SNR: *t*(101.62) = -3.63, *p*_corr_ < 0.001).

There was no main effect of age (*F*(1,50) = 0.92, *p* = 0.34, η = 0.014) and no significant interaction of age with intelligibility or predictability (Figure 2D).

### fMRI Results – Task-related activity

#### Intelligibility effects in bilateral superior temporal cortex are independent of age

First, we were interested in task-related activity sensitive to increasingly intelligible speech, irrespective of its predictability. A conjunction analysis revealed increased activity in bilateral primary auditory regions in the superior temporal gyri for increasingly intelligible speech comprehension independent of age (Supplementary Figure 2A and Supplementary Table 3). The direct comparison between age groups revealed increased activation in left and right precentral gyrus as well as left and right cerebellum in older participants. Contrary, older participants exhibited less activation in medial frontal regions than young listeners.

#### Semantic regions are commonly recruited when speech is highly predictable

Next, we explored brain regions that support the processing of semantic information, irrespective of intelligibility. Here we identified that activation common to both age groups during highly predictable speech was located in left AG, pMTG and precuneus/paracingulate gyrus (Supplementary Figure 2B and Supplementary Table 4). These regions are considered key regions (pMTG and AG) or extended regions (precuneus) of the semantic system in young and older adults (Binder et al., 2009; Hoffman & Morcom, 2018). Direct comparisons between age groups revealed increased activation in a large cluster in the precuneus in young compared to older participants during the processing of highly predictable speech. No other differences were found. The reversed contrast revealed that the pre-SMA was the only region commonly recruited by both groups for low predictable sentences irrespective of intelligibility (Supplementary Figure 2C).

#### The recruitment of semantic versus cingulo-opercular regions depends on the predictability of speech when intelligibility is increasing but not on age

Finally, we were interested in brain regions that are differentially affected by predictability when intelligibility is varied parametrically (i.e., the interaction between intelligibility and predictability). To this end, we modelled intelligibility as linear parametric modulation of high and low predictable sentences. In both groups, we found higher activation for high versus low predictable sentences under increasing intelligibility in several large left- and right-hemispheric clusters (Figure 3). Overall, the activation pattern was broadly similar in both age groups, while cluster sizes were considerably larger in young adults (Figure 3). The conjunction analysis of the interaction contrast revealed that left angular gyrus (PGa & PGp), right angular gyrus (PGp) as well as left posterior middle temporal gyrus were recruited by both age groups (Figure 3A, left, Supplementary Table 5). Note that, opposed to the main effect of predictability, the homologous right angular gyrus seems to play a role in both age groups when sentence intelligibility is varied. Young adults showed increased activity in a cluster comprising the postcentral gyrus in the right hemisphere. No other significant differences between age groups were found.

**Figure 3.**
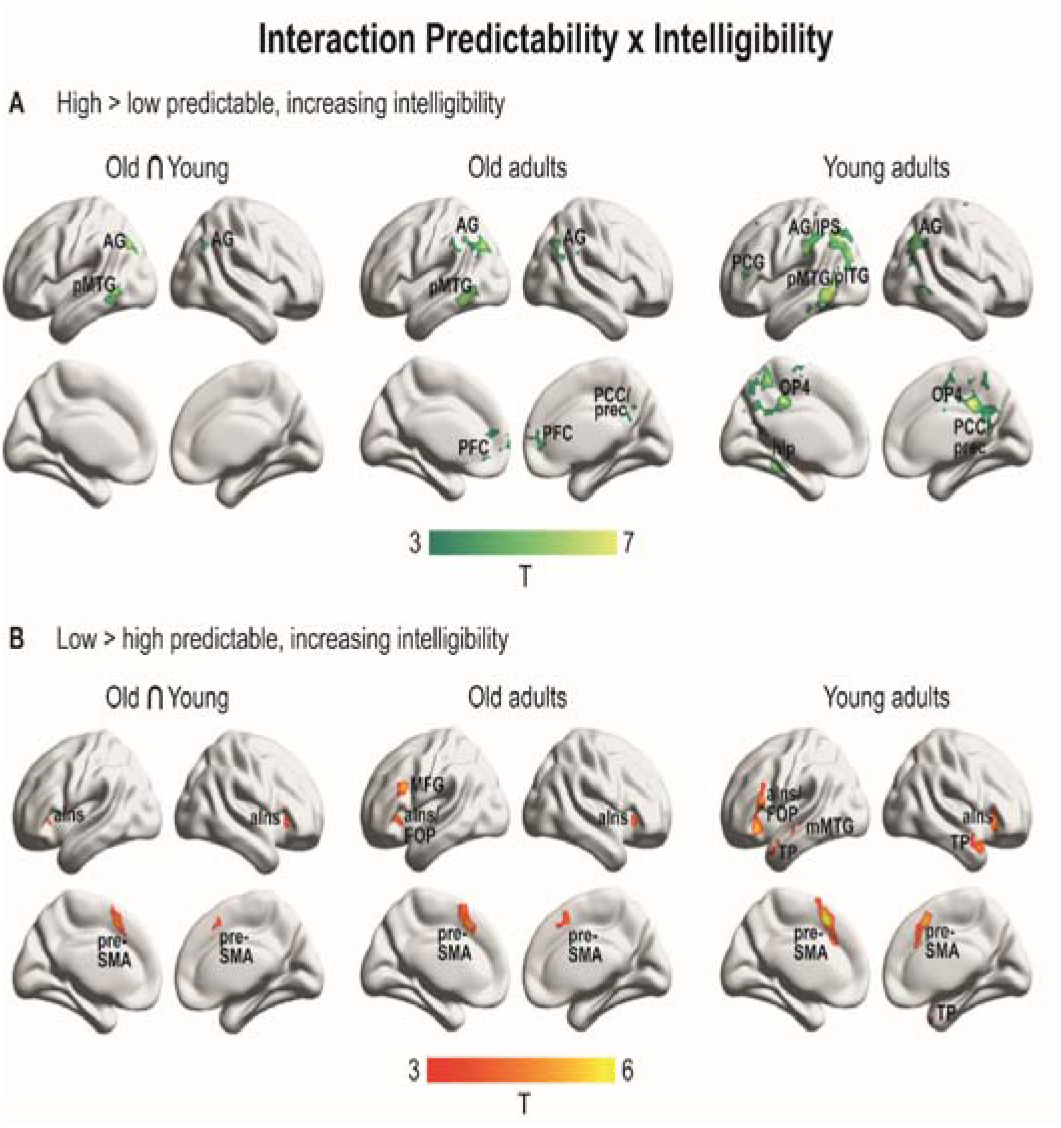
fMRI results showing the interaction effect of intelligibility and predictability. **A** Brain regions that show significant effects for the interaction contrast (high x increasing intelligibility > low x increasing intelligibility). The left column shows regions commonly activated by both groups. Activation maps of the interaction contrast are displayed separately for old (middle) and young participants (right) for visualization. **B** Brain regions that show significant effects in the opposite direction of the interaction (low x increasing intelligibility > high x increasing intelligibility). All activation maps are thresholded at cluster-level *p*_*FWE*_ < 0.05, with a voxel-wise threshold of *p* < 0.001, AG = angular gyrus, pMTG/pITG = posterior middle/inferior temporal gyrus, PFC = prefrontal cortex, PCC/prec = posterior cingulate cortex/precuneus, PCG = precentral gyrus, IPS = inferior parietal sulcus, OP4 = parietal operculum, hip = hippocampus, prec = precuneus, FOP = frontal operculum, aIns = anterior insula, pre-SMA = pre-supplementary motor area, TP = temporal pole, mMTG = mid-middle temporal gyrus.

The opposite direction of the interaction contrast targeted brain regions sensitive to sentences of low predictability under increasing intelligibility. There were no significant differences between age groups. Both groups showed significant activity in the pre-SMA and bilateral anterior insulae, extending into the frontal operculum (Figure 3B, Supplementary Table 6). These regions are associated with the domain-general cingulo-opercular network (Dosenbach et al., 2008). Note that in contrast to the main effect of predictability reported above, older adults here indeed showed increased activity in the anterior insular regions. The clusters in pre-SMA and bilateral insulae were identified as common activation across both age groups via a conjunction analysis and were used as seed regions for the effective connectivity analysis, together with left pMTG and bilateral AG.

### fMRI Results – Effective connectivity

At the network level, we investigated commonalities and age-related differences in effective connectivity driven by the interactive effect of predictability and intelligibility within the semantic and cingulo-opercular network and between both networks. We followed a dual-step procedure comparing (1) age-related differences in the network architecture of the cingulo-opercular network based on previous findings (Rysop et al. 2021) and (2) age-related differences in a larger network configuration containing seed regions based on the exact coordinates from the univariate conjunction analyses reported above (Figure 4A).

**Figure 4.**
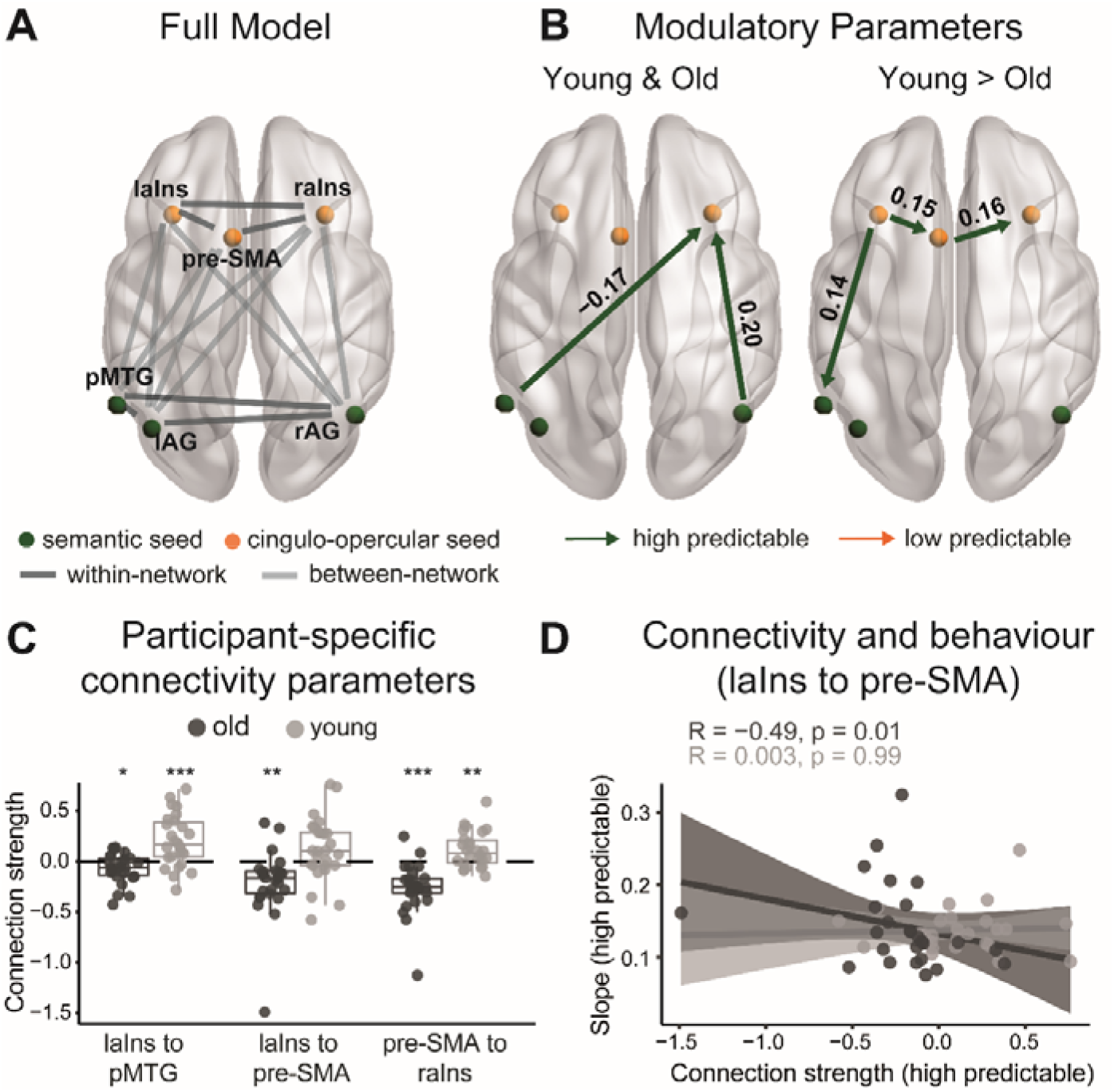
DCM PEB results. **A** Illustration of the full DCM model; semantic seeds are coloured in green, cingulo-opercular seeds are coloured in orange, within-network connections are shown in dark grey, between-network connections are shown in light grey; laIns/raIns = left/right anterior insula, pMTG = posterior middle temporal gyrus, l/rAG = left/right angular gyrus, pre-SMA = pre-supplementary motor area. **B** Group results of the Bayesian model average (BMA) of the modulatory parameters (B-parameters), thresholded at a posterior probability of > 0.95. Modulatory effects of high (green arrow) and low (orange arrow) predictable sentences with parametrically modulated intelligibility. Left: Modulatory effects of the intelligibility x predictability interaction that are common for both groups. Right: Age-related differences (young > old adults) for the modulatory effects of high predictable sentences. **C** Individual modulatory parameters for high predictable sentences at each connection showing an age-related difference. Asterisks indicate connections that significantly differ from zero, * = *p* < 0.05, ** = *p* < 0.01, *** = *p* < 0.001. **D** Negative correlation between connectivity (modulation from left anterior insula to pre-SMA) and the slope parameter from the psychometric curve for high predictable sentences, shaded area represents 95 % confidence intervals.

First, we asked whether the cingulo-opercular model that was identified as the best model explaining the data of young participants would also succeed in explaining the data of older participants. Overall, the exceedance probabilities for all families of models included in the DCM analysis were remarkably low in the older subgroup, with the highest exceedance probability of 0.35 (opposed to 0.73 in young participants). No clear winning family of models could be identified. The winning model that was identified in young participants corresponded to the second probable model in the older group, with an exceedance probability of 0.24 (see Supplementary Figure 3) and an average of 14.36 % explained variance (opposed to 22.13 % in young participants), thus indicating that the winning model of young participants must not necessarily serve as an optimal model for older participants as well. Independent-samples t-tests conducted on the parameter estimates that were extracted from the modulated connections of that model (i.e., winning model in young participants) indicated no significant age-related differences (see Supplementary Figure 3).

#### High predictability modulates connectivity from semantic regions to the right anterior insula in both age groups

As a next step, we conducted a PEB DCM analysis that is more powerful in the analysis of age-related differences in effective connectivity as it allows a larger network space encompassing seed regions from both networks of interest across both age groups. The full model yielded an average of 22.78 % explained variance across both groups (22.75 % for old participants, 22.82 % for young participants). First, we examined which connections were modulated by either high or low predictable sentences under increasing intelligibility in both groups (e.g., commonalities across participants). Here we found that high predictability had an inhibitory influence on the connection from left pMTG to right aIns (Figure 4B, left; see also Table 1 and Supplementary Figure 5 for individual parameters of common modulatory effects). There was no intrinsic connection between these regions, thus these regions form a functional interaction during the task, but do not interact with each other during rest (see Supplementary Tables 7 and 8 and Supplementary Figure 4 for intrinsic connections). Moreover, high predictability had an excitatory influence on the connection from right AG to right aIns, further strengthening the positive intrinsic connection between these regions. Low predictable sentences did not have a significant effect on any of the connections, aside from weak evidence for the inhibitory influence from pMTG to right aIns (posterior probability = 0.64).

**Table 1.** Modulatory parameter estimates from the Bayesian model average (BMA) reflecting commonalities in connectivity across groups.

| Commonalities across age groups |  |  |  |  |
| --- | --- | --- | --- | --- |
| Connection | Modulatory connectivity |  |  |  |
|  | High * SNR | Pp | Low * SNR | Pp |
| *Between Network Connections* |  |  |  |  |
| rAG → ralns | **0.20** | **1** | 0.00 | 0 |
| pMTG → ralns | **-0.17** | **1** | -0.09 | 0.64 |

Note: Only connections with at least one parameter exceeding a posterior probability (Pp) > .95 are shown and highlighted in bold. See Supplementary Table 7 for commonalities in intrinsic connectivity and self-connections.

#### Young adults exhibit stronger connectivity within cingulo-opercular regions and between left insula and pMTG when sentences are highly predictable and increasingly intelligible

Next, we looked at connections that showed age-related differences in the modulatory effect of high and low predictable sentences. Age-related differences were found in the modulatory influence of high predictability on three connections: from left aIns to pMTG, from left aIns to pre-SMA and from pre-SMA to right aIns (Figure 4B, right; see also Table 2). Coupling between these connections was stronger for younger adults. Testing connectivity values for each group against zero, revealed that the modulatory effect was negative (i.e., inhibitory) in older adults for all three connections (*p* < 0.05; see Figure 4C for individual parameters of connections that showed age-related differences and Supplementary Figure 5 for individual parameters of connections with a similar modulatory effects). In contrast, all modulatory parameters of young participants were positive (i.e., excitatory), except for the non-significant modulation of the connectivity from left anterior insula to pre-SMA (*p* = 0.06). None of the connections had a stronger coupling for older compared to younger adults. Low predictable sentences did not have a significant modulatory influence on any connection.

**Table 2.** Modulatory parameter estimates from the Bayesian model average (BMA) reflecting differences in connectivity between groups.

| Differences between age groups (young > old) |  |  |  |  |
| --- | --- | --- | --- | --- |
| Connection | Modulatory connectivity |  |  |  |
|  | High * SNR | Pp | Low * SNR | Pp |
| *Within Cingulo-Opercular Network* |  |  |  |  |
| lalns → pre-SMA | **0.15** | **1** | 0.00 | 0.00 |
| pre-SMA → ralns | **0.16** | **1** | 0.00 | 0.00 |
| *Between Network Connections* |  |  |  |  |
| lalns → pMTG | **0.14** | **1** | 0.00 | 0.00 |

Note: Only connections with at least one parameter exceeding a posterior probability (Pp) > .95 are shown and highlighted in bold. For the differences between age groups positive values indicate stronger connectivity for young adults. See Supplementary Table 8 for differences in intrinsic connectivity.

To test the predictive validity of the age-related connectivity differences, we performed a leave-one-out cross-validation on the three connectivity parameters that differed between age groups. The out-of-samples Pearson correlation between the predicted and actual group membership was significant for all three connections (left aIns → pMTG: *r* = 0.29, *p* = 0.028; left aIns → pre-SMA: *r* = 0.28, *p* = 0.023; pre-SMA → right aIns: *r* = 0.34, *p* = 0.006). Thus, the effect sizes were large enough to predict a left-out participant’s age group based on their connectivity.

Finally, we were interested whether age differences in connectivity strength were associated with behavioural performance. To this end, we correlated connectivity parameters that showed significant group effects with the respective parameters from the psychometric curves, representing the behavioural predictability gain (threshold and slope parameter). We found a significant negative relationship between the slope parameter of high predictable sentences (representing the sensitivity to changes in intelligibility) and the coupling strength for the connection from left anterior insula to the pre-SMA (i.e., *within* the cingulo-opercular network) in old participants (*rho* = -0.49, *p* = 0.012; Figure 4D). Those older participants with a more negative connectivity were more sensitive to changes in intelligibility of highly predictable sentences at the behavioural level. Put differently, increased inhibition between these two domain-general regions was associated with higher accuracy at the behavioural level in old participants. This connectivity-behaviour association was not significant in young participants (*rho* = 0.046, *p* = 0.83). No other connectivity-behaviour relationships were significant.

Together, these results demonstrate that connectivity within the semantic and the cingulo-opercular network, as well as between both networks, was mainly modulated by high predictable sentences. Younger participants exhibited stronger coupling between subregions of the cingulo-opercular network and between left anterior insula and pMTG when speech was increasingly intelligible and highly predictive compared to their older counterparts. Decreased connectivity within the cingulo-opercular network was associated with increased task performance selectively in older adults.

## Discussion

In this study, we investigated commonalities and differences during speech comprehension under challenging listening conditions in young and middle-aged to old participants. First, we found that both age groups showed a comparable behavioural predictability gain, reflected in overlapping psychometric curves. Second, depending on the semantic context of increasingly intelligible speech, both groups recruited distributed left-hemispheric semantic and cingulo-opercular brain regions. Third, young and old listeners differed in effective connectivity during highly predictable and intelligible speech: Young adults exhibited overall stronger connectivity between regions of the cingulo-opercular network and between left anterior insula and pMTG. Moreover, these interactions were excitatory in young adults but inhibitory in old adults. Finally, the degree of the inhibitory influence from left anterior insula to pre-SMA was predictive of the behavioural sensitivity towards changes in intelligibility for high predictable sentences in older adults only. Our results demonstrate that the predictability gain is relatively preserved in middle-aged to older adults with good peripheral hearing when stimulus intelligibility is adjusted to individual hearing abilities. Accordingly, age-related differences at the neural level are subtle and manifest in effective connectivity, where inhibition between cingulo-opercular regions is associated with better comprehension when speech is highly predictive.

### Old listeners benefit from semantic context as much as young listeners

Our experiment was designed to assure comparable task difficulty for young and old listeners, as older listeners are known to have increased difficulties with hearing speech in challenging listening conditions even if they have relatively normal audiograms (Pichora-Fuller et al., 2017). Although we only included older participants with relatively good peripheral hearing as measured by an audiogram, they had significantly higher speech reception thresholds than young listeners. This apparent mismatch converges with the notion that hearing acuity in quiet (as measured by pure-tone audiometry) is not a good predictor of hearing abilities in noise (Schneider et al., 2010). Moreover, this stresses the importance of taking individual differences in hearing abilities into account when conducting cross-sectional experiments in the auditory domain, for instance by carefully adjusting stimulus material.

Indeed, when controlling for differences in peripheral hearing abilities, both age groups showed a comparable benefit from semantic context, suggesting that younger and older listeners did not differ in their ability to exploit contextual cues. In earlier studies, older adults showed a similar or even slightly enhanced benefit from semantic context (e.g., Pichora-Fuller, 2008; Pichora-Fuller et al., 1995). This was explained by a stronger recruitment of top-down contextual information to compensate for the age-related decline in bottom-up sensory information in older adults. If this was the case, we would have expected a relative “overuse” of semantic cues when auditory degradation was adjusted to individual hearing acuity, manifesting in a larger predictability gain in older adults. However, we did not observe age-related differences at the behavioural level. In general, our finding that older adults show a comparable beneficial effect from contextual cues is consistent with previous findings that reported a comparable predictability gain for older listeners across a wide range of intelligibility manipulations (e.g., Dubno et al., 2000; Goy et al., 2013; Humes, 1996; Sheldon et al., 2008).

### Age does not affect which brain regions are recruited in speech comprehension

At the level of task-related brain activity, both age groups showed speech-sensitive engagement of left and right auditory regions, encompassing Heschl’s gyrus and the superior temporal cortex, as well as the right parietal operculum. The common recruitment of bilateral primary auditory regions is well in line with the literature, where hypoactivation of the primary auditory cortex was observed in listeners with age-related hearing loss (Peelle et al., 2011) or listeners with normal hearing but worse task performance (Wong et al., 2009). With respect to age-related differences, older adults exhibited less activity in frontal regions, including right temporal pole and dorsal anterior cingulate cortex, which may reflect unspecific dedifferentiation effects related to ageing. In contrast, stronger activity in the bilateral precentral gyrus and the cerebellum for older relative to younger participants may reflect a different listening strategy such as inner rehearsal in older participants (Guediche, Holt, Laurent, Lim, & Fiez, 2015). Indeed, a stronger contribution of the cerebellum in older adults has been previously reported during picture naming and was linked to articulatory and phonological processes (Chen & Desmond, 2005; Ferré, Jarret, Brambati, Bellec, & Joanette, 2020).

With respect to the interaction of predictability and intelligibility, older participants showed broadly the same activation pattern as young adults, with generally smaller cluster sizes. Common activity across age groups was mainly found in left angular gyrus and posterior middle temporal gyrus regions associated with semantic (control) processes (Binder et al., 2009; Jackson, 2021). This is further in line with a meta-analysis (Hoffman & Morcom, 2018) reporting overall similar activation for young and old participants in semantic tasks. In that meta-analysis, differences between age groups manifested as additional right-hemispheric activity in older adults and were observed when task performance of older participants was poor. We found no evidence for such additional domain-general activity in older adults, most likely because we assured that the difficulty of the sentence repetition task was comparable across participants. On the contrary, young listeners showed stronger activity in an additional brain region, namely the precuneus, when contrasting high and low predictable sentences irrespective of intelligibility. Increased activity in the precuneus/posterior cingulate is often observed in semantic tasks and may serve as an interface between semantic and memory systems (Binder et al., 2009). The exact role of the precuneus/posterior cingulate complex in semantic cognition is not clear, as these regions are associated with numerous cognitive functions (see Jung & Lambon Ralph, 2021; Smallwood et al., 2021). Age-related differences in this region might reflect a consequence of neural dedifferentiation, manifesting as a change in the relative balance of this region.

Aside from the expected association of left angular gyrus activity with the predictability gain, we also found increased activity in right angular gyrus. The effective connectivity between right angular gyrus and right anterior insula was strengthened in young and old listeners when speech was highly predictable and increasingly intelligible. These findings point towards a potential demand-related role of right angular gyrus in semantic processing. There is mounting evidence that both left and right angular gyri are involved in (combinatorial) semantic processes (Binder et al., 2009; Golestani et al., 2013; Graessner, Zaccarella, & Hartwigsen, 2021; Graves, Binder, Desai, Conant, & Seidenberg, 2010; Price, Bonner, Peelle, & Grossman, 2015). During speech comprehension under challenging listening conditions, right angular gyrus showed increased activity for related words at higher intelligibility in young participants, but at a lower threshold than left angular gyrus (Golestani et al., 2013). This supports the notion that right angular gyrus might indeed be specifically recruited during semantic processing when task demands increase.

When semantic context was low, both age groups showed increased activity in regions of the cingulo-opercular network for more intelligible speech, which was expected based on the literature (Peelle, 2018; Vaden Jr. et al., 2015, 2013) and likely helps to maintain stable performance under increased task demands (Peelle, 2018). However, contrary to our hypothesis, we did not observe a stronger recruitment of cingulo-opercular regions in older adults, which has been reported in previous work (Reuter-Lorenz & Cappell, 2008), and was linked to successful performance in older listeners (Erb & Obleser, 2013). The difference between the previous and present work is likely explained by our adaptive tracking paradigm which calibrated noise levels to the individual hearing abilities. Our results imply that previously reported differences between age groups may have resulted from increased task demands for older participants due to differences in peripheral hearing.

### Age affects connectivity within and between networks recruited in degraded speech comprehension

Finally, at the neural network level, we found common interaction patterns across groups but also age-related differences in effective connectivity. In both groups, task-related changes in connectivity were observed selectively for high predictable sentences (with linearly increasing intelligibility), but not for low predictable sentences. Independent of age, high predictable sentences increased the inhibitory influence of left pMTG on the right anterior insula, as well as the excitatory influence of right AG on the right anterior insula. This finding points towards a complex interaction pattern of semantic-specific and domain-general areas during speech comprehension under challenging listening conditions even when task-relevant semantic information is available.

Although we did not find age-related differences at the level of activation, effective connectivity patterns within the cingulo-opercular network (left anterior insula to pre-SMA, pre-SMA to right anterior insula), and between cingulo-opercular and semantic regions (left anterior insula to left pMTG) differed as a function of age. Connectivity between these areas was overall stronger for young adults. Notably, the direction of the influence differed between both groups, with excitatory connectivity in young and inhibitory connectivity in old listeners. This pattern of overall reduced within-network connectivity in cognitive control networks in the ageing brain converges with several previous findings, although it remains debated whether these changes reflect processes of decline or compensation (Campbell, Grady, Ng, & Hasher, 2012; Madden et al., 2010; O’Connell & Basak, 2018 but see Grady et al., 2010). In our study, the degree of the inhibitory influence between cingulo-opercular regions was associated with a steeper slope of the psychometric curve for high predictable speech in older adults only. This reflects higher sensitivity towards intelligibility in highly predictable speech selectively in older adults and suggests a potential beneficial effect of inhibition within the cingulo-opercular network when semantic context can be used to predict upcoming speech. Further evidence supporting this notion comes from a recent study demonstrating that increased connectivity within domain-general networks was associated with less efficient behavioural performance in a semantic task in older adults (Martin, Saur, & Hartwigsen, 2021). Importantly, the opposite pattern was observed for young adults, which converges with our finding of increased connectivity within the cingulo-opercular network in young adults. Together, the present results show that such inhibition of the interactions in the domain-general cingulo-opercular network may constitute a key mechanism underlying the predictability gain in the ageing brain.

It is noteworthy however, that younger listeners exhibited a pattern of inhibition within the cingulo-opercular network in a previous investigation (Rysop et al., 2021). In the present study, the same model appeared less suitable for older adults, as revealed by a low amount of explained variance captured by the model alongside low exceedance probability. This apparently conflicting finding demonstrates that effective connectivity results must be interpreted with caution and with respect to the broader context of its setup. As the PEB DCM approach yielded higher levels of explained variance, considers the (un)certainty of participants’ parameters and allows a larger model space enabling the investigation of within and between network interactions, we believe that this approach is more suitable to investigate age-related differences in effective connectivity.

### Limitations

To our knowledge, this study represents the first investigation of age-related differences in the effective connectivity underlying the predictability gain during speech processing under challenging listening conditions. However, one drawback of the DCM methodology is that it only allows to include a limited number of seed regions. Therefore, conclusions about interactions with other regions and functional networks cannot be drawn based on our analysis. Future studies may address how the presence of semantic context alters network configurations between functional networks, such as the default mode, cingulo-opercular or the fronto-parietal control network on a larger level, and how such configurations change with age.

### Conclusions

The present study shows that young and old adults recruit a similar set of semantic and cingulo-opercular brain regions during speech comprehension under challenging listening conditions when intelligibility was calibrated to the individual hearing thresholds. This highlights the importance of carefully adjusting stimulus intensities to individual hearing levels in ageing research. With such precautions, age-related differences in task-related recruitment are rather subtle and manifest only at the network level. Older adults showed overall decreased connectivity between semantic and cingulo-opercular regions as well as within the cingulo-opercular network. Nevertheless, the degree of inhibition within the cingulo-opercular network was positively related to speech comprehension in older adults only, demonstrating the behavioural relevance of such interactions. These findings provide new insight into age-related changes in the network configuration underlying the predictability gain.

## Author contributions

**Anna Uta Rysop**: Investigation, Methodology, Data Curation, Formal analysis, Software, Validation, Visualization, Writing – Original Draft, Writing – Review & Editing. **Lea-Maria Schmitt**: Methodology, Software, Formal analysis, Writing – Review & Editing. **Jonas Obleser**: Conceptualization, Funding Acquisition, Writing – Review & Editing, Resources, Supervision. **Gesa Hartwigsen**: Conceptualization, Funding Acquisition, Writing – Review & Editing, Resources, Supervision.

## Acknowledgements

The authors would like to thank Isabel Gebhardt and Rebekka Luckner for assistance in data acquisition. We also thank Anke Kummer, Sylvie Neubert and Manuela Hofmann for assistance during the fMRI measurements. We are grateful to Julia Erb for providing the stimulus materials and to the University of Minnesota Center for Magnetic Resonance Imaging for providing the fMRI multiband sequence software.

## Transparency

We report how we determined our sample size, all data exclusions, all inclusion/exclusion criteria, whether inclusion/exclusion criteria were established prior to data analysis, all manipulations, and all measures in the study. No parts of the study procedures and analyses were pre-registered in a time-stamped, institutional registry prior to the research being conducted.

Raw neuroimaging data are protected under the General Data Protection Regulation (EU) and can only be made available from the corresponding authors upon reasonable request. The written version of the experimental stimuli (German SPIN sentences) can be found in the appendix of Erb et al., 2012. Readers seeking access to the auditory stimuli and custom analysis code that was used in addition to the analysis softwares SPM and JASP should contact the corresponding authors.

## References

Adank, P. (2012). The neural bases of difficult speech comprehension and speech production: Two Activation Likelihood Estimation (ALE) meta-analyses. Brain and Language, 122(1), 42–54. https://doi.org/10.1016/j.bandl.2012.04.014

Alain, C., Du, Y., Bernstein, L. J., Barten, T., & Banai, K. (2018). Listening under difficult conditions□: An activation likelihood estimation meta-analysis. Human Brain Mapping, 39(February), 2695–2709. https://doi.org/10.1002/hbm.24031

Aschenbrenner, S., Tucha, O., & Lange, K. W. (2000). Regensburger Wortflüssigkeits-Test: RWT. Hogrefe, Verlag für Psychologie.

Binder, J. R., Desai, R. H., Graves, W. W., & Conant, L. L. (2009). Where is the semantic system? A critical review and meta-analysis of 120 functional neuroimaging studies. Cerebral Cortex, 19(12), 2767–2796. https://doi.org/10.1093/cercor/bhp055

Cameron, A. C., & Windmeijer, F. A. G. (1997). An R-squared measure of goodness of fit for some common nonlinear regression models. Journal of Econometrics, 77(2), 329–342. https://doi.org/10.1016/S0304-4076(96)01818-0

Campbell, K. L., Grady, C. L., Ng, C., & Hasher, L. (2012). Age differences in the frontoparietal cognitive control network: Implications for distractibility. Neuropsychologia, 50(9), 2212–2223. https://doi.org/10.1016/j.neuropsychologia.2012.05.025

Chen, S. H. A., & Desmond, J. E. (2005). Temporal dynamics of cerebro-cerebellar network recruitment during a cognitive task. Neuropsychologia, 43(9), 1227–1237. https://doi.org/10.1016/J.NEUROPSYCHOLOGIA.2004.12.015

Darwin, C. J. (2008). Listening to speech in the presence of other sounds. Philosophical Transactions of the Royal Society B: Biological Sciences, 363(1493), 1011–1021. https://doi.org/10.1098/rstb.2007.2156

Desjardins, J. L., & Doherty, K. A. (2013). Age-Related Changes in Listening Effort for Various Types of Masker Noises. Ear & Hearing, 34(3), 261–272. https://doi.org/10.1097/AUD.0b013e31826d0ba4

Dijkstra, N., Zeidman, P., Ondobaka, S., Van Gerven, M. A. J., & Friston, K. (2017). Distinct Top-down and Bottom-up Brain Connectivity during Visual Perception and Imagery. Scientific Reports, 7(1), 1–9. https://doi.org/10.1038/s41598-017-05888-8

Dosenbach, N. U. F., Fair, D. A., Cohen, A. L., Schlaggar, B. L., & Petersen, S. E. (2008). A dual-networks architecture of top-down control. Trends in Cognitive Sciences, 12(3), 99–105. https://doi.org/10.1016/j.tics.2008.01.001

Dubno, J. R., Ahlstrom, J. B., & Horwitz, A. R. (2000). Use of context by young and aged adults with normal hearing. The Journal of the Acoustical Society of America, 107(1), 538–546. https://doi.org/10.1121/1.428322

Eckert, M. A., Walczak, A., Ahlstrom, J., Denslow, S., Horwitz, A., & Dubno, J. R. (2008). Age-related effects on word recognition: Reliance on cognitive control systems with structural declines in speech-responsive cortex. JARO - Journal of the Association for Research in Otolaryngology, 9(2), 252–259. https://doi.org/10.1007/s10162-008-0113-3

Eickhoff, S. B., Paus, T., Caspers, S., Grosbras, M.-H., Evans, A. C., Zilles, K., & Amunts, K. (2007). Assignment of functional activations to probabilistic cytoarchitectonic areas revisited. NeuroImage, 36(3), 511–521. https://doi.org/10.1016/j.neuroimage.2007.03.060

Eickhoff, S. B., Stephan, K. E., Mohlberg, H., Grefkes, C., Fink, G. R., Amunts, K., & Zilles, K. (2005). A new SPM toolbox for combining probabilistic cytoarchitectonic maps and functional imaging data. NeuroImage, 25(4), 1325–1335. https://doi.org/10.1016/j.neuroimage.2004.12.034

Erb, J., Henry, M. J., Eisner, F., & Obleser, J. (2012). Auditory skills and brain morphology predict individual differences in adaptation to degraded speech. Neuropsychologia, 50(9), 2154–2164. https://doi.org/10.1016/j.neuropsychologia.2012.05.013

Erb, J., & Obleser, J. (2013). Upregulation of cognitive control networks in older adults’ speech comprehension. Frontiers in Systems Neuroscience, 7(December), 1–13. https://doi.org/10.3389/fnsys.2013.00116

Federmeier, K. D. (2007). Thinking ahead: The role and roots of prediction in language comprehension. Psychophysiology, 44(4), 491–505. https://doi.org/10.1111/j.1469-8986.2007.00531.x

Feinberg, D. A., Moeller, S., Smith, S. M., Auerbach, E., Ramanna, S., Glasser, M. F., … Yacoub, E. (2010). Multiplexed echo planar imaging for sub-second whole brain fmri and fast diffusion imaging. PLoS ONE, 5(12). https://doi.org/10.1371/journal.pone.0015710

Ferré, P., Jarret, J., Brambati, S. M., Bellec, P., & Joanette, Y. (2020). Task-Induced Functional Connectivity of Picture Naming in Healthy Aging: The Impacts of Age and Task Complexity. Neurobiology of Language, 1(2), 161–184. https://doi.org/10.1162/nol_a_00007

Fitzhugh, M. C., Braden, B. B., Sabbagh, M. N., Rogalsky, C., & Baxter, L. C. (2019). Age-Related Atrophy and Compensatory Neural Networks in Reading Comprehension. Journal of the International Neuropsychological Society, 25(6), 569–582. https://doi.org/10.1017/S1355617719000274

Fitzhugh, M. C., Schaefer, S. Y., Baxter, L. C., & Rogalsky, C. (2021). Cognitive and neural predictors of speech comprehension in noisy backgrounds in older adults. Language, Cognition and Neuroscience, 36(3), 269– 287. https://doi.org/10.1080/23273798.2020.1828946

Folstein, M. F., Folstein, S. E., & McHugh, P. R. (1975). “Mini-mental state.” Journal of Psychiatric Research, 12(3), 189–198. https://doi.org/10.1016/0022-3956(75)90026-6

Friston, K. J., Harrison, L., & Penny, W. (2003). Dynamic causal modelling. NeuroImage, 19(4), 1273–1302. https://doi.org/10.1016/S1053-8119(03)00202-7

Friston, K. J., Litvak, V., Oswal, A., Razi, A., Stephan, K. E., van Wijk, B. C. M., … Zeidman, P. (2016). Bayesian model reduction and empirical Bayes for group (DCM) studies. NeuroImage, 128, 413–431. https://doi.org/10.1016/j.neuroimage.2015.11.015

Friston, K. J., Worsley, K. J., Frackowiak, R. S. J., Mazziotta, J. C., & Evans, A. C. (1994). Assessing the significance of focal activations using their spatial extent. Human Brain Mapping, 1(3), 210–220. https://doi.org/10.1002/hbm.460010306

Fründ, I., Haenel, N. V., & Wichmann, F. A. (2011). Inference for psychometric functions in the presence of nonstationary behavior. Journal of Vision, 11(6), 16–16. https://doi.org/10.1167/11.6.16

Geerligs, L., Renken, R. J., Saliasi, E., Maurits, N. M., & Lorist, M. M. (2015). A Brain-Wide Study of Age-Related Changes in Functional Connectivity. Cerebral Cortex, 25(July), 1987–1999. https://doi.org/10.1093/cercor/bhu012

Golestani, N., Hervais-Adelman, A., Obleser, J., & Scott, S. K. (2013). Semantic versus perceptual interactions in neural processing of speech-in-noise. NeuroImage, 79, 52–61. https://doi.org/10.1016/j.neuroimage.2013.04.049

Gordon-Salant, S. (2005). Hearing loss and aging: New research findings and clinical implications. Journal of Rehabilitation Research and Development, 42(4 SUPPL. 2), 9–24. https://doi.org/10.1682/JRRD.2005.01.0006

Gordon-Salant, S., & Fitzgibbons, P. J. (1995). Recognition of Multiply Degraded Speech by Young and Elderly Listeners. Journal of Speech, Language, and Hearing Research, 38(5), 1150–1156. https://doi.org/10.1044/jshr.3805.1150

Goy, H., Pelletier, M., Coletta, M., & Kathleen Pichora-Fuller, M. (2013). The effects of semantic context and the type and amount of acoustic distortion on lexical decision by younger and older adults. Journal of Speech, Language, and Hearing Research, 56(6), 1715–1732. https://doi.org/10.1044/1092-4388(2013/12-0053)

Grady, C. L., Protzner, A. B., Kovacevic, N., Strother, S. C., Afshin-Pour, B., Wojtowicz, M., … McIntosh, A. R. (2010). A Multivariate Analysis of Age-Related Differences in Default Mode and Task-Positive Networks across Multiple Cognitive Domains. Cerebral Cortex, 20(6), 1432–1447. https://doi.org/10.1093/cercor/bhp207

Graessner, A., Zaccarella, E., & Hartwigsen, G. (2021). Differential contributions of left-hemispheric language regions to basic semantic composition. Brain Structure and Function, 226(2), 501–518. https://doi.org/10.1007/s00429-020-02196-2

Graves, W. W., Binder, J. R., Desai, R. H., Conant, L. L., & Seidenberg, M. S. (2010). Neural correlates of implicit and explicit combinatorial semantic processing. NeuroImage, 53(2), 638–646. https://doi.org/10.1016/j.neuroimage.2010.06.055

Guediche, S., Holt, L. L., Laurent, P., Lim, S., & Fiez, J. A. (2015). Evidence for Cerebellar Contributions to Adaptive Plasticity in Speech Perception. Cerebral Cortex, 25(July), 1867–1877. https://doi.org/10.1093/cercor/bht428

Halai, A. D., Welbourne, S. R., Embleton, K., & Parkes, L. M. (2014). A comparison of dual gradient-echo and spin-echo fMRI of the inferior temporal lobe. Human Brain Mapping, 35(8), 4118–4128. https://doi.org/10.1002/hbm.22463

Hartwigsen, G., Golombek, T., & Obleser, J. (2015). Repetitive transcranial magnetic stimulation over left angular gyrus modulates the predictability gain in degraded speech comprehension. Cortex, 68, 100–110. https://doi.org/10.1016/j.cortex.2014.08.027

Hoffman, P., & Morcom, A. M. (2018). Age-related changes in the neural networks supporting semantic cognition: A meta-analysis of 47 functional neuroimaging studies. Neuroscience and Biobehavioral Reviews, 84(November 2017), 134–150. https://doi.org/10.1016/j.neubiorev.2017.11.010

Humes, L. E. (1996). Speech understanding in the elderly. Journal of American Audiology, 7(3), 161–167.

Jackson, R. L. (2021). The neural correlates of semantic control revisited. NeuroImage, 224(September 2020), 117444. https://doi.org/10.1016/j.neuroimage.2020.117444

Jefferies, E. (2013). The neural basis of semantic cognition: Converging evidence from neuropsychology, neuroimaging and TMS. Cortex. https://doi.org/10.1016/j.cortex.2012.10.008

Jung, J., & Lambon Ralph, M. A. (2021). Distinct but cooperating brain networks supporting semantic cognition. BioRxiv, (July). https://doi.org/10.1101/2021.07.19.452716

Kalikow, D. N., Stevens, K. N., & Elliott, L. L. (1977). Development of a test of speech intelligibility in noise using sentence materials with controlled word predictability. The Journal of the Acoustical Society of America, 61(5), 1337–1351. https://doi.org/10.1121/1.381436

Kass, R. E., & Raftery, A. E. (1995). Bayes Factors. Journal of the American Statistical Association, 90(430), 773–795. https://doi.org/10.1080/01621459.1995.10476572

Kollmeier, B., Gilkey, R. H., & Sieben, U. K. (1988). Adaptive staircase techniques in psychoacoustics: A comparison of human data and a mathematical model. The Journal of the Acoustical Society of America, 83(5), 1852–1862. https://doi.org/10.1121/1.396521

Li, S.-C., Lindenberger, U., & Sikström, S. (2001). Aging cognition: from neuromodulation to representation. Trends in Cognitive Sciences, 5(11), 479–486. https://doi.org/10.1016/S1364-6613(00)01769-1

Madden, D. J., Costello, M. C., Dennis, N. A., Davis, S. W., Shepler, A. M., Spaniol, J., … Cabeza, R. (2010). Adult age differences in functional connectivity during executive control. NeuroImage, 52(2), 643–657. https://doi.org/10.1016/j.neuroimage.2010.04.249

Marreiros, A. C., Kiebel, S. J., & Friston, K. J. (2008). Dynamic causal modelling for fMRI: A two-state model. NeuroImage, 39(1), 269–278. https://doi.org/10.1016/j.neuroimage.2007.08.019

Martin, S., Saur, D., & Hartwigsen, G. (2021). Age-dependent contribution of domain-general networks to semantic cognition. Cerebral Cortex, 2020.11.06.371153. Retrieved from https://doi.org/10.1101/2020.11.06.371153

Miller, G. A. (1947). The masking of speech. Psychological Bulletin, 44(2), 105–129. https://doi.org/10.1037/h0055960

Nichols, T., Brett, M., Andersson, J., Wager, T., & Poline, J.-B. (2005). Valid conjunction inference with the minimum statistic. NeuroImage, 25(3), 653–660. https://doi.org/10.1016/j.neuroimage.2004.12.005

O’Connell, M. A., & Basak, C. (2018). Effects of task complexity and age-differences on task-related functional connectivity of attentional networks. Neuropsychologia, 114, 50–64. https://doi.org/10.1016/j.neuropsychologia.2018.04.013

Obleser, J., & Kotz, S. A. (2010). Expectancy constraints in degraded speech modulate the language comprehension network. Cerebral Cortex, 20(3), 633–640. https://doi.org/10.1093/cercor/bhp128

Obleser, J., Wise, R. J. S., Dresner, A. M., & Scott, S. K. (2007). Functional Integration across Brain Regions Improves Speech Perception under Adverse Listening Conditions. Journal of Neuroscience, 27(9), 2283– 2289. https://doi.org/10.1523/JNEUROSCI.4663-06.2007

Oldfield, R. C. (1971). The assessment of handedness: The Edinburgh inventory. Neuropsychologia.

Payne, B. R., & Federmeier, K. D. (2018). Contextual constraints on lexico-semantic processing in aging: Evidence from single-word event-related brain potentials. Brain Research, 1687, 117–128. https://doi.org/10.1016/j.brainres.2018.02.021

Peelle, J. E. (2018). Listening Effort: How the Cognitive Consequences of Acoustic Challenge Are Reflected in Brain and Behavior. Ear and Hearing, 39(2), 204–214. https://doi.org/10.1097/AUD.0000000000000494

Peelle, J. E., Troiani, V., Grossman, M., & Wingfield, A. (2011). Hearing Loss in Older Adults Affects Neural Systems Supporting Speech Comprehension. The Journal of Neuroscience, 31(35), 12638–12643. https://doi.org/10.1523/JNEUROSCI.2559-11.2011

Penny, W., Stephan, K. E., Daunizeau, J., Rosa, M. J., Friston, K. J., Schofield, T. M., & Leff, A. P. (2010). Comparing families of dynamic causal models. PLoS Computational Biology, 6(3). https://doi.org/10.1371/journal.pcbi.1000709

Pichora-Fuller, M. K. (2008). Use of supportive context by younger and older adult listeners: Balancing bottom-up and top-down information processing. International Journal of Audiology, 47(SUPPL. 2). https://doi.org/10.1080/14992020802307404

Pichora-Fuller, M. K., Alain, C., & Schneider, B. A. (2017). Older Adults at the Cocktail Party. In J. C.. Middlebrooks, J. Z.. Simon, A. N.. Popper, & R. R. Fay (Eds.), The Auditory System at the Cocktail Party (1st ed., pp. 227–259). Springer. https://doi.org/10.1007/978-3-319-51662-2_9

Pichora-Fuller, M. K., Schneider, B. A., & Daneman, M. (1995). How young and old adults listen to and remember speech in noise. The Journal of the Acoustical Society of America, 97(1), 593–608. https://doi.org/10.1121/1.412282

Poldrack, R. A., Nichols, T., & Mumford, J. (2011). Handbook of Functional MRI Data Analysis. Cambridge: Cambridge University Press. https://doi.org/10.1017/CBO9780511895029

Price, A. R., Bonner, M. F., Peelle, J. E., & Grossman, M. (2015). Converging evidence for the neuroanatomic basis of combinatorial semantics in the angular gyrus. Journal of Neuroscience, 35(7), 3276–3284. https://doi.org/10.1523/JNEUROSCI.3446-14.2015

R, C. T. (2021). R: A Language and environment for statistical computing. Vienna, Austria: R Foundation for Statistical Computing. Retrieved from https://www.r-project.org/

Reitan, R. M. (1958). Validity of the Trail Making Test as an Indicator of Organic Brain Damage. Perceptual and Motor Skills, 8(3), 271–276. https://doi.org/10.2466/pms.1958.8.3.271

Reuter-Lorenz, P. A., & Cappell, K. A. (2008). Neurocognitive Aging and the Compensation Hypothesis. Current Directions in Psychological Science, 17(3), 177–182. https://doi.org/10.1111/j.1467-8721.2008.00570.x

Rogers, C. S., & Peelle, J. E. (2022). Interactions Between Audition and Cognition in Hearing Loss and Aging. In L. L. Holt, J. E. Peelle, A. B. Coffin, A. N. Popper, & R. R. Fay (Eds.), Speech Perception (pp. 227–252). Springer, ASA Press. https://doi.org/10.1007/978-3-030-81542-4_9

Rysop, A. U., Schmitt, L. M., Obleser, J., & Hartwigsen, G. (2021). Neural modelling of the semantic predictability gain under challenging listening conditions. Human Brain Mapping, 42(1), 110–127. https://doi.org/10.1002/hbm.25208

Sala-Llonch, R., Bartres-Faz, D., & Junque, C. (2015). Reorganization of brain networks in aging: a review of functional connectivity studies. Frontiers in Psychology, 6(May), 1–11. https://doi.org/10.3389/fpsyg.2015.00663

Salthouse, T. A., Atkinson, T. M., & Berish, D. E. (2003). Executive Functioning as a Potential Mediator of Age-Related Cognitive Decline in Normal Adults. Journal of Experimental Psychology: General, 132(4), 566– 594. https://doi.org/10.1037/0096-3445.132.4.566

Schneider, B. A., Pichora-Fuller, M. K., & Daneman, M. (2010). The effects of senescant changes in audition and cognition on spoken language comprehension. In S. Gordon-Salant, R. D. Frisina, A. N. Popper, & R. R. Fay (Eds.), Springer Handbook of Auditory Research: The Aging Auditory System: Perceptual Characterizsation and Neural Bases of Presbycusis (pp. 167–210).

Schütt, H. H., Harmeling, S., Macke, J. H., & Wichmann, F. A. (2016). Painfree and accurate Bayesian estimation of psychometric functions for (potentially) overdispersed data. Vision Research, 122, 105–123. https://doi.org/10.1016/j.visres.2016.02.002

Seghier, M. L. (2013). The Angular Gyrus. The Neuroscientist, 19(1), 43–61. https://doi.org/10.1177/1073858412440596

Setton, R., Mwilambwe-Tshilobo, L., Girn, M., Lockrow, A. W., Baracchini, G., Hughes, C., … Spreng, R. N. (2022). Age differences in the functional architecture of the human brain. Cerebral Cortex, 1–21. https://doi.org/10.1093/cercor/bhac056

Shannon, R. V., Zeng, F.-G., Kamath, V., Wygonski, J., & Ekelid, M. (1995). Speech Recognition with Primarily Temporal Cues. Science, 270(5234), 303–304. https://doi.org/10.1126/science.270.5234.303

Sheldon, S., Pichora-Fuller, M. K., & Schneider, B. A. (2008). Priming and sentence context support listening to noise-vocoded speech by younger and older adults. The Journal of the Acoustical Society of America, 123(1), 489–499. https://doi.org/10.1121/1.2783762

Siegel, J. S., Power, J. D., Dubis, J. W., Vogel, A. C., Church, J. A., Schlaggar, B. L., & Petersen, S. E. (2014). Statistical improvements in functional magnetic resonance imaging analyses produced by censoring high-motion data points. Human Brain Mapping, 35(5), 1981–1996. https://doi.org/10.1002/hbm.22307

Smallwood, J., Bernhardt, B. C., Leech, R., Bzdok, D., Jefferies, E., & Margulies, D. S. (2021). The default mode network in cognition: a topographical perspective. Nature Reviews Neuroscience, 22(8), 503–513. https://doi.org/10.1038/s41583-021-00474-4

Stephan, K. E., Penny, W., Daunizeau, J., Moran, R. J., & Friston, K. J. (2009). Bayesian model selection for group studies. NeuroImage, 46(4), 1004–1017. https://doi.org/10.1016/j.neuroimage.2009.03.025

Uddin, L. Q., Yeo, B. T. T., & Spreng, R. N. (2019). Towards a Universal Taxonomy of Macro-scale Functional Human Brain Networks. Brain Topography, (0123456789). https://doi.org/10.1007/s10548-019-00744-6

Vaden Jr., K. I., Kuchinsky, S. E., Ahlstrom, J., Dubno, J. R., & Eckert, M. A. (2015). Cortical Activity Predicts Which Older Adults Recognize Speech in Noise and When. Journal of Neuroscience, 35(9), 3929–3937. https://doi.org/10.1523/JNEUROSCI.2908-14.2015

Vaden Jr., K. I., Kuchinsky, S. E., Cute, S. L., Ahlstrom, J., Dubno, J. R., & Eckert, M. A. (2013). The Cingulo-Opercular Network Provides Word-Recognition Benefit. Journal of Neuroscience, 33(48), 18979–18986. https://doi.org/10.1523/JNEUROSCI.1417-13.2013

Verhaeghen, P. (2003). Aging and vocabulary score: A meta-analysis. Psychology and Aging, 18(2), 332–339. https://doi.org/10.1037/0882-7974.18.2.332

Wagenmakers, E.-J., Love, J., Marsman, M., Jamil, T., Ly, A., Verhagen, J., … Morey, R. D. (2018). Bayesian inference for psychology. Part II: Example applications with JASP. Psychonomic Bulletin & Review, 25(1), 58–76. https://doi.org/10.3758/s13423-017-1323-7

Wechsler, D. (1955). Manual for the Wechsler Adult Intelligence Scale. Manual for the Wechsler Adult Intelligence Scale. Oxford, England: Psychological Corp.

Wichmann, F. A., & Hill, N. J. (2001). Wichmann psychometric and Hill I: fitting, sampling, and goodness–of–fit. Perception & Psychophysics, 63(8), 1293--1313.

Wilsch, A., Henry, M. J., Herrmann, B., Maess, B., & Obleser, J. (2015). Alpha Oscillatory Dynamics Index Temporal Expectation Benefits in Working Memory. Cerebral Cortex, 25(7), 1938–1946. https://doi.org/10.1093/cercor/bhu004

Wingfield, A., Amichetti, N. M., & Lash, A. (2015). Cognitive aging and hearing acuity: Modeling spoken language comprehension. Frontiers in Psychology, 6(MAY), 1–13. https://doi.org/10.3389/fpsyg.2015.00684

Wingfield, A., & Stine-Morrow, E. A. L. (2000). Language and Speech. In F. I. M. Craik & T. A. Salthouse (Eds.), The handbook of aging and cognition (pp. 359–416).

Wlotko, E. W., Federmeier, K. D., & Kutas, M. (2012). To predict or not to predict: Age-related differences in the use of sentential context. Psychology and Aging, 27(4), 975–988. https://doi.org/10.1037/a0029206

Wong, P. C. M., Jin, J. X., Gunasekera, G. M., Abel, R., Lee, E. R., & Dhar, S. (2009). Aging and cortical mechanisms of speech perception in noise. Neuropsychologia, 47(3), 693–703. https://doi.org/10.1016/j.neuropsychologia.2008.11.032

Xia, M., Wang, J., & He, Y. (2013). BrainNet Viewer: A Network Visualization Tool for Human Brain Connectomics. PLoS ONE, 8(7), e68910. https://doi.org/10.1371/journal.pone.0068910

Zeidman, P., Jafarian, A., Corbin, N., Seghier, M. L., Razi, A., Price, C. J., & Friston, K. J. (2019). A guide to group effective connectivity analysis, part 1: First level analysis with DCM for fMRI. NeuroImage, 200(June), 174–190. https://doi.org/10.1016/j.neuroimage.2019.06.031

Zeidman, P., Jafarian, A., Seghier, M. L., Litvak, V., Cagnan, H., Price, C. J., & Friston, K. J. (2019). A guide to group effective connectivity analysis, part 2: Second level analysis with PEB. NeuroImage, 200(June), 12– 25. https://doi.org/10.1016/j.neuroimage.2019.06.032

Zhang, H., Gertel, V. H., Cosgrove, A. L., & Diaz, M. T. (2021). Age-related differences in resting-state and task-based network characteristics and cognition: a lifespan sample. Neurobiology of Aging, 101, 262–272. https://doi.org/10.1016/j.neurobiolaging.2020.10.025

Zonneveld, H. I., Pruim, R. H., Bos, D., Vrooman, H. A., Muetzel, R. L., Hofman, A., … Vernooij, M. W. (2019). Patterns of functional connectivity in an aging population: The Rotterdam Study. NeuroImage, 189, 432– 444. https://doi.org/10.1016/j.neuroimage.2019.01.041

