## Supplementary Material for "Age-related differences in the neural network interactions underlying the predictability gain"

**Supplementary Methods**

**Behavioural Analyses**

We fitted cumulative Gaussian sigmoid functions as beta-binomial observer models. The beta-binomial model extends the standard binomial model by an overdispersion parameter which allows to carry out statistical inference on data affected by performance fluctuations challenging the serial independence of trials ((Schütt, Harmeling, Macke, & Wichmann, 2016). Goodness of psychometric fits was assessed by comparing the empirical deviance (i.e., two times the log-likelihood ratio of the saturated model to the fitted model) with a Monte Carlo simulated deviance distribution (Wichmann & Hill, 2001). For each psychometric curve, a reference distribution was created by (a) randomly drawing from a binomial distribution with n = 18 trials and p = fitted probability of a correct response for each intelligibility level, (b) calculating the deviance across intelligibility levels, and (c) repeating this procedure for 10,000 samples.

**Dynamic Causal Modelling (DCM) - Standard DCM Setup**

For the standard DCM we used a design matrix with one regressor containing the onsets of all experimental stimuli as driving input. Modulatory parameters were incorporated as six regressors encoding high and low predictability at paired intelligibility levels of low (–9 and –4 dB SNR relative to SRT), medium (–1 and +1 dB SNR relative to SRT) and high intelligibility (+4 and +9 dB SNR relative to SRT; see Rysop et al., 2021 for a detailed description of this DCM setup).

At the first-level, we specified 84 models per participant. All models had full bidirectional intrinsic connections and inhibitory self-connections. The models differed in the configuration of the driving input (7 variants) and the modulatory input (12 variants). The driving input was set to either one, two or all regions at a time. The modulatory influence was set to either one connection or two afferent or efferent connections at a time. To reduce the model space to a reasonable number, models were grouped in 12 families of models defined by the modulatory input (Penny et al., 2010). Within each family of models the modulatory input was kept constant while the driving input was varied.

At the second-level we identified the most probable family of models given the data using the random-effects Bayesian Model Selection procedure (Penny et al., 2010; Stephan, Penny, Daunizeau, Moran, & Friston, 2009). In this procedure, each family of models is assigned an exceedance probability that allows to quantify how good the model describes the underlying data. We focussed on the question whether the winning family of models, that we identified in Rysop et al., 2021, would also succeed in explaining the data of the older subgroup. We further extracted the percentage of explained variance to obtain a parameter of the model fit for older participants. Finally, we extracted single-participant modulatory parameter estimates from the winning model of the previous publication (Rysop et al., 2021). Parameter estimates were submitted to an independent-samples t-test to investigate age-related differences on modulated connections between regions. Independent-samples t-tests were calculated in JASP (version 0.9.1), parameter estimates were visualized with R (version 4.0.2).

**Table 1.** Demographic variables and results of neuropsychological testing of older participants (n=26)

| **Demographics** |  |  |
| --- | --- | --- |
| Age (years) | 62.31 (6.17) |  |
| Gender (F:M) | 16:10 |  |
| Education (years) | 17.44 (3.23) |  |
| Pure Tone Average dB (left ear; right ear) | 18.48 (4.82); 17.62 (6.15) |  |
| **Neuropsychological Tests** | **Raw Score** | **Interpretation** |
| Mini Mental State Examination (max. 30) | 28.92 (1.86) | no cognitive impairment |
| Digit Span Task forward | 8.38 (1.62) |  |
| Digit Span Task backward | 6.69 (1.86) |  |
| Trail Making Test A (time in s) | 26.28 (9.05) | perfectly normal |
| Trail Making Test B (time in s) | 55.85 (16.65) | perfectly normal |
| Word Fluency (sum *food*) | 35.62 (8.93) | PR = 62 |
| Word Fluency (sum *fruits & sports*) | 21.38 (5.53) | PR = 53 |

Note: Values represent the mean, standard deviation is denoted in brackets, PR = percentile rank

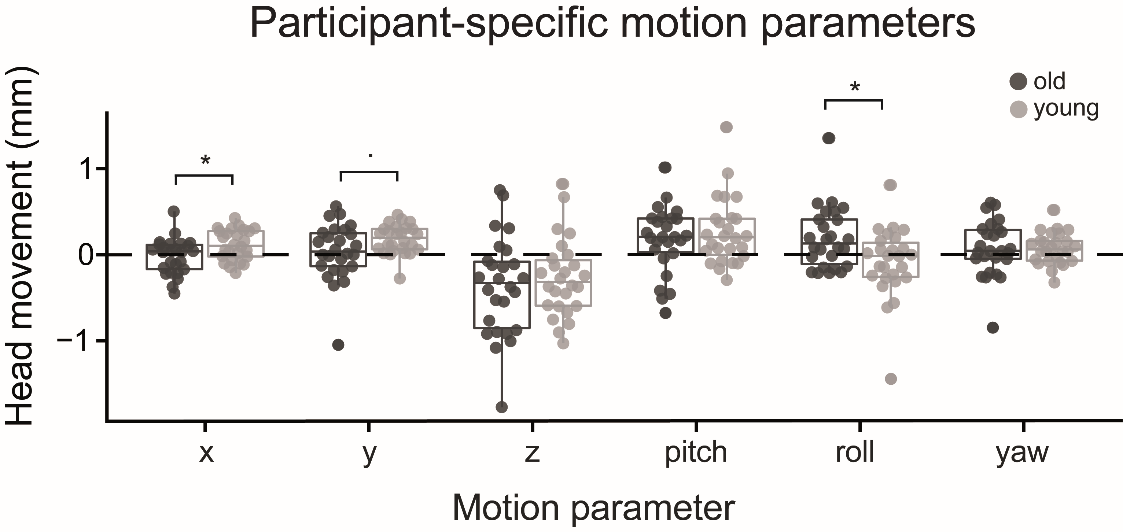

**Figure 1.** Overview of the translational and rotational motion parameters obtained from the rigid-body transformation of the realignment step for young and old participants. Each dot represents the participant-specific average of the respective parameter. * = *p* < 0.05, *.* = *p* = 0.05.

**Table 2.** Statistical Analysis of the parameters derived from the psychometric curves.

| Parameter | Effect | F | *p* | η² |
| --- | --- | --- | --- | --- |
| Threshold | Age | 0.498 | 0.484 | 0.009 |
|  | **Predictability** | **302.426** | **<0.001** | **0.414** |
|  | Age * predictability | 2.075 | 0.156 | 0.005 |
| Slope | Age | 0.181 | 0.672 | 0.002 |
|  | **Predictability** | **7.455** | **0.009** | **0.076** |
|  | Age * predictability | 0.870 | 0.355 | 0.010 |

Note: Significant effects are highlighted in bold

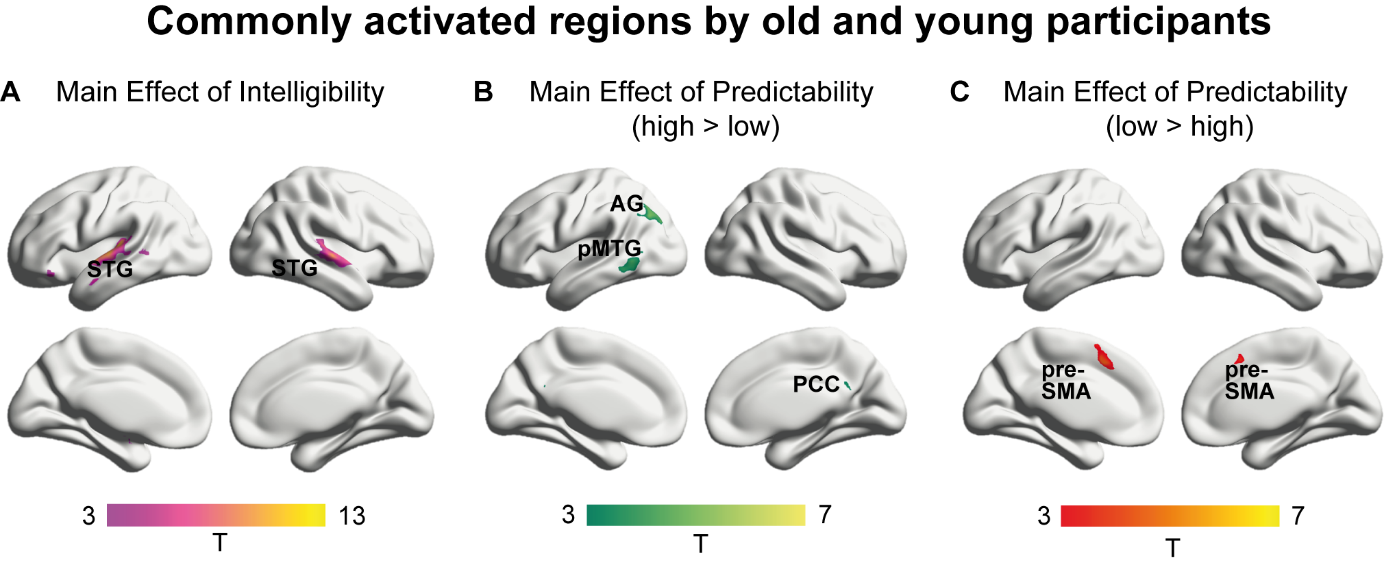

**Figure 2**. **Activation maps showing the results of the conjunction analysis**. Brain regions displayed are recruited by both age groups. **A** Brain regions sensitive to increasingly intelligible speech irrespective of predictability, thresholded at peak-level *p_FWE_* < 0.05. **B** Brain regions sensitive to predictability of speech irrespective of intelligibility for high predictable sentences, thresholded at a voxel-wise *p* < 0.001 and a cluster-wise *p_FWE_* < 0.05. **C** Brain regions sensitive to predictability of speech irrespective of intelligibility for low predictable sentences, thresholded at a voxel-wise *p* < 0.001 and a cluster-wise *p_FWE_* < 0.05. STG = superior temporal gyrus, AG = angular gyrus, pMTG = posterior middle temporal gyrus, PCC/prec = posterior cingulate cortex/precuneus, pre-SMA = pre-supplementary motor area.

**Table 3.** fMRI results. Main Effect of Intelligibility.

| **Location** | **Hem** | **MNI** | | | ***t*** | **Cluster Size** |
| --- | --- | --- | --- | --- | --- | --- |
|  |  | x | y | z |  |  |
| ***Old ∩ Young*** | | | | | | |
| Superior temporal gyrus (Area TE 3) | L | -62 | -10 | 2 | 7.99 | 243 |
| Superior temporal gyrus (Area TE 1.0) | L | -44 | -24 | 10 | 7.55 | Subcluster |
| Superior temporal gyrus (Area TE 1.1) | L | -40 | -32 | 10 | 7.49 | Subcluster |
| Parietal operculum (Area OP4) | R | 60 | -4 | 2 | 7.51 | 125 |
| Superior temporal gyrus (Area TE 1.0) | R | 46 | -20 | 8 | 7.17 | Subcluster |
| Superior temporal gyrus (Area TE 3) | R | 63 | -12 | 5 | 7.03 | Subcluster |
| ****Old > Young*** | | | | | | |
| Precentral gyrus (Area 3b) | L | -62 | -4 | 18 | 6.40 | 277 |
| Precentral gyrus (Area 3a) | L | -40 | -17 | 35 | 5.73 | Subcluster |
| Precentral gyrus (Area 3a) | L | -50 | -10 | 25 | 5.47 | Subcluster |
| Precentral gyrus (Area 3a) | R | 48 | -7 | 28 | 5.86 | 276 |
| Precentral gyrus (Area 1) | R | 53 | -7 | 48 | 5.21 | Subcluster |
| Cerebellum | L | -14 | -62 | -22 | 5.50 | 279 |
| Cerebellum | R | 18 | -37 | -30 | 18 | 63 |
| ****Young > Old*** | | | | | | |
| Frontal pole (Area Fo1) | R | 10 | 38 | -20 | 4.50 | 95 |
| Dorsal anterior cingulate (Area s32) | R | 0 | 40 | -18 | 4.25 | Subcluster |
| Dorsal anterior cingulate (Area s32) | L | -2 | 26 | -20 | 3.96 | Subcluster |

Results are thresholded at a *p* < 0.05 (FWE peak-level correction). Contrasts marked with an asterisk are significant at *p* < 0.05 (FWE cluster-level correction).

**Table 4.** fMRI results. Main Effect of Predictability.

| **Location** | **Hem** | **MNI** | | | ***t*** | **Cluster Size** |
| --- | --- | --- | --- | --- | --- | --- |
| **High > Low predictable sentences** |  | x | y | z |  |  |
| ***Old ∩ Young*** | | | | | | |
| Angular gyrus (Area PGp) | L | -40 | -67 | 35 | 6.02 | 200 |
| Angular gyrus (Area PGa) | L | -42 | -70 | 42 | 5.99 | Subcluster |
| Angular gyrus (Area PGp) | L | -40 | -80 | 30 | 5.45 | Subcluster |
| Posterior middle temporal gyrus | L | -60 | -60 | -5 | 5.19 | 196 |
| Posterior middle temporal gyrus | L | -60 | -50 | -10 | 4.62 | Subcluster |
| Posterior middle temporal gyrus | L | -52 | -64 | 5 | 4.16 | Subcluster |
| Posterior cingulate gyrus | R | 8 | -52 | 25 | 4.24 | 49 |
| Precuneus | L | -7 | -52 | 25 | 3.94 | Subcluster |
| Precuneus | L | -12 | -60 | 12 | 3.87 | Subcluster |
| ***Young > Old*** | | | | | | |
| Precuneus | L | -20 | -52 | 20 | 4.98 | 55 |
| Precuneus | L | -14 | -44 | 28 | 4.53 | Subcluster |
| **Low > high predictable sentences** |  |  |  |  |  |  |
| ***Old ∩ Young*** |  |  |  |  |  |  |
| Pre-SMA (Area 6mr) | L | -7 | 13 | 48 | 5.40 | 217 |
| Pre-SMA (Area 6mr) | L | -4 | 10 | 60 | 4.65 | Subcluster |
| Pre-SMA (Area 6mr) | R | 6 | 16 | 48 | 4.61 | Subcluster |

Results are thresholded at voxel-wise *p* < 0.001 and a cluster-wise *p* < 0.05; FWE-corrected.

**Table 5.** fMRI results. High > low predictable sentences with increasing intelligibility.

| **Location** | **Hem** | **MNI** | | | ***t*** | **Cluster Size** |
| --- | --- | --- | --- | --- | --- | --- |
|  |  | x | y | z |  |  |
| ***Old participants*** | | | | | | |
| Inferior parietal lobe (Area PGp) | L | -42 | -70 | 35 | 6.17 | 196 |
| Inferior parietal lobe (Area PGa) | L | -42 | -70 | 42 | 5.65 | Subcluster |
| Inferior parietal lobe (Area PFm) | L | -54 | -54 | 40 | 4.43 | Subcluster |
| Posterior middle temporal gyrus | L | -57 | -62 | -2 | 5.85 | 297 |
| Posterior inferior temporal gyrus | L | -60 | -50 | -12 | 5.04 | Subcluster |
| Posterior inferior temporal gyrus | L | -52 | -54 | -12 | 4.99 | Subcluster |
| Paracingulate gyrus (Area p24c) | R | 10 | 46 | 5 | 5.84 | 622 |
| Accumbens (Area 25) | L | -10 | 10 | -10 | 5.10 | Subcluster |
| Frontal pole (Area p32) | R | 6 | 60 | 20 | 5.06 | Subcluster |
| Lateral occipital cortex (Area hOc4la) | R | 58 | -62 | 0 | 5.14 | 66 |
| Inferior parietal lobe (Area PGp) | R | 46 | -64 | 35 | 5.04 | 66 |
| Precuneus | L | -7 | -70 | 22 | 4.46 | 56 |
| Posterior cingulate gyrus | R | 8 | -52 | 25 | 3.77 | Subcluster |
| Precuneus | R | 6 | -60 | 22 | 3.76 | Subcluster |
| Superior parietal lobe (Area 5Ci) | L | -12 | -34 | 40 | 4.28 | 48 |
| Precuneus | L | -2 | -30 | 38 | 3.99 | Subcluster |
| Precuneus | L | -4 | -40 | 35 | 3.68 | Subcluster |
| ***Young participants*** | | | | | | |
| Posterior cingulate gyrus | L | -2 | -37 | 40 | 8.74 | 5100 |
| Inferior parietal sulcus (Area hIP5) | L | -34 | -80 | 30 | 8.38 | Subcluster |
| Superior parietal lobe (Area 5Ci) | R | 6 | -32 | 38 | 8.33 | Subcluster |
| Inferior parietal lobe (Area PGp) | R | 46 | -67 | 42 | 7.59 | 693 |
| Inferior parietal lobe (Area PGa) | R | 48 | -60 | 35 | 6.71 | Subcluster |
| Inferior parietal lobe (Area PGa) | R | 48 | -70 | 32 | 6.28 | Subcluster |
| Hippocampus (DG) | L | -27 | -27 | -12 | 5.89 | 178 |
| Hippocampus (CA1) | L | -30 | -40 | -18 | 5.78 | Subcluster |
| Hippocampus (CA1) | L | -34 | -30 | -15 | 5.71 | Subcluster |
| Accumbens (Area 35) | L | -10 | 10 | -10 | 5.51 | 46 |
| Insular cortex | L | -37 | -10 | -8 | 5.26 | 83 |
| Putamen | L | -30 | -4 | -10 | 4.99 | Subcluster |
| Insular Cortex | L | -37 | 0 | -10 | 4.54 | Subcluster |
| Putamen | R | 28 | 6 | 8 | 5.13 | 99 |
| Putamen | R | 30 | -12 | 2 | 4.70 | Subcluster |
| Putamen | R | 30 | -2 | 5 | 4.34 | Subcluster |
| Parietal Operculum (Area OP4) | R | 56 | -4 | 10 | 4.97 | 52 |
| Parietal Operculum (Area OP4) | R | 60 | -14 | 2 | 4.08 | Subcluster |
| Parietal Operculum (Area OP1) | R | 60 | -14 | 12 | 3.48 | Subcluster |
| Posterior middle temporal gyrus | R | 58 | -52 | 0 | 4.86 | 155 |
| Posterior middle temporal gyrus | R | 63 | -44 | -5 | 4.36 | Subcluster |
| Posterior middle temporal gyrus | R | 48 | -54 | 10 | 3.88 | Subcluster |
| Temporal occipital fusiform cortex | R | 40 | -50 | -12 | 4.86 | 82 |
| Superior temporal gyrus (Area TE 1.1) | R | 38 | -32 | 15 | 4.68 | 96 |
| Inferior parietal lobe (Area PFcm) | R | 56 | -27 | 10 | 4.48 | Subcluster |
| Inferior parietal lobe (Area PFcm) | R | 60 | -32 | 15 | 4.32 | Subcluster |
| Insular Cortex | L | -37 | -4 | 12 | 4.55 | 55 |
| Insular Cortex | L | -34 | 3 | 10 | 4.47 | Subcluster |
| Precentral Gyrus | L | -44 | 0 | 20 | 4.06 | Subcluster |
| ***Old ∩ Young*** | | | | | | |
| Posterior middle temporal gyrus | L | -57 | -60 | -2 | 5.80 | 244 |
| Posterior middle temporal gyrus | L | -57 | -54 | -10 | 4.49 | Subcluster |
| Posterior middle temporal gyrus | L | -64 | -47 | -5 | 4.41 | Subcluster |
| Inferior parietal lobe (Area PGp) | L | -42 | -70 | 35 | 5.79 | 148 |
| Inferior parietal lobe (Area PGp) | L | -40 | -80 | 30 | 5.50 | Subcluster |
| Inferior parietal lobe (Area PGa) | L | -52 | -60 | 38 | 3.90 | Subcluster |
| Inferior parietal lobe (Area PGp) | R | 46 | -64 | 35 | 5.04 | 65 |
| ***Young > Old*** | | | | | | |
| Postcentral gyrus (Area 2) | R | 18 | -40 | 60 | 4.74 | 51 |
| Postcentral gyrus (Area 2) | R | 3 | -40 | 58 | 4.29 | Subcluster |

Results are thresholded at voxel-wise *p* < 0.001 and a cluster-wise *p* < 0.05; FWE-corrected.

**Table 6.** fMRI results. Low > high predictable sentences with increasing intelligibility.

| **Location** | **Hem** | **MNI** | | | ***t*** | **Cluster Size** |
| --- | --- | --- | --- | --- | --- | --- |
|  |  | x | y | z |  |  |
| ***Old participants*** | | | | | | |
| Inferior frontal gyrus (Area 44) | L | -40 | 16 | 22 | 5.61 | 144 |
| Pre-SMA (Area 6mr) | L | -7 | 13 | 48 | 5.33 | 349 |
| Pre-SMA (Area 6mr) | L | -2 | 23 | 42 | 5.19 | Subcluster |
| Pre-SMA (Area 6mr) | L | -4 | 10 | 60 | 4.96 | Subcluster |
| Anterior insula (Area Id7) | R | 33 | 23 | 2 | 5.08 | 70 |
| Anterior insula (Area Id7) | R | 28 | 30 | -5 | 4.80 | Subcluster |
| Middle frontal gyrus | L | -34 | 3 | 48 | 4.75 | 51 |
| Anterior insula (Area Id7) | L | -30 | 23 | -2 | 4.74 | 71 |
| Frontal operculum (Area OP8) | L | -42 | 23 | -2 | 3.80 | Subcluster |
| ***Young participants*** | | | | | | |
| Pre-SMA (Area 6mr) | L/R | 0 | 16 | 52 | 7.23 | 514 |
| Pre-SMA (Area 6mr) | R | 6 | 26 | 35 | 6.35 | Subcluster |
| Pre-SMA (Area 6mr) | L | -7 | 16 | 48 | 6.23 | Subcluster |
| Frontal operculum (Area OP9) | L | -50 | 23 | 2 | 6.65 | 482 |
| Frontal operculum (Area OP8) | L | -42 | 26 | -10 | 6.07 | Subcluster |
| Anterior insula (Area Id7) | L | -30 | 26 | -2 | 5.37 | Subcluster |
| Anterior insula (Area Id7) | R | 33 | 28 | -5 | 6.17 | 160 |
| Anterior insula (Area Id7) | R | 30 | 20 | 5 | 4.36 | Subcluster |
| Inferior frontal gyrus (Area 45) | R | 46 | 20 | 0 | 4.06 | Subcluster |
| Temporal pole | R | 36 | 0 | -25 | 5.39 | 259 |
| Temporal pole | R | 53 | 6 | -25 | 5.10 | Subcluster |
| Anterior middle temporal gyrus | R | 46 | -7 | -25 | 5.00 | Subcluster |
| Middle middle temporal gyrus | L | -52 | -14 | -15 | 5.29 | 57 |
| ***Old ∩ Young*** | | | | | | |
| Pre-SMA (Area 6mr) | L | -7 | 13 | 48 | 5.33 | 282 |
| Pre-SMA (Area 6mr) | L | -2 | 10 | 60 | 4.85 | Subcluster |
| Pre-SMA (Area 6mr) | R | 6 | 26 | 35 | 4.65 | Subcluster |
| Anterior insula (Area Id7) | R | 33 | 23 | 0 | 4.78 | 57 |
| Anterior insula (Area Id7) | R | 30 | 30 | -5 | 4.23 | Subcluster |
| Anterior insula (Area Id7) | L | -30 | 23 | -2 | 4.74 | 60 |
| Frontal operculum (Area OP8) | L | -42 | 23 | -2 | 3.80 | Subcluster |

Results are thresholded at voxel-wise *p* < 0.001 and a cluster-wise *p* < 0.05; FWE-corrected.

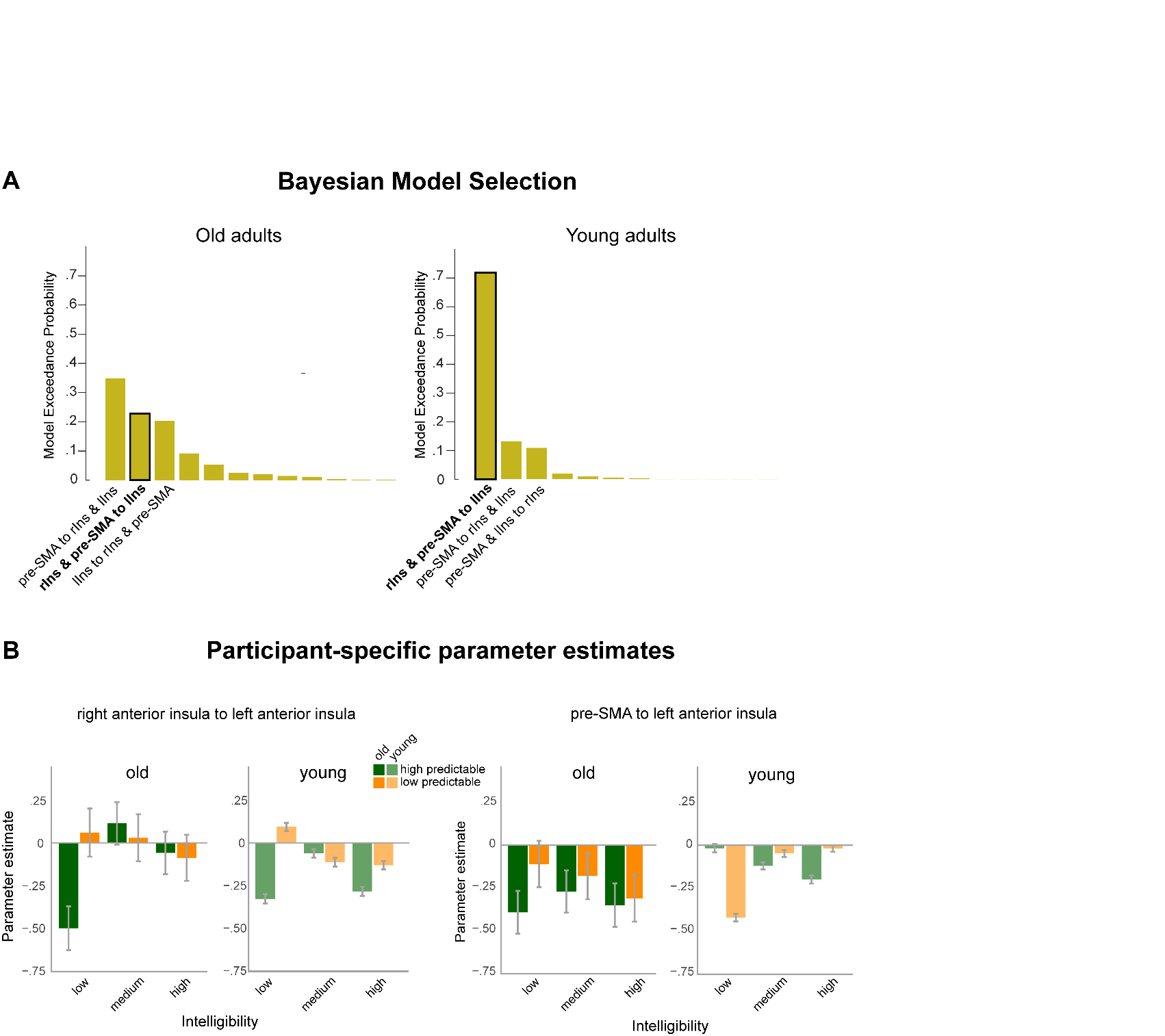

**Figure 3. Standard DCM Results**. **A** Results of the Bayesian Model Selection (BMS) of the DCM within the cingulo-opercular network in old (left) and young adults (right). Bar height represents the exceedance probability of a certain family of models, names are given for the three most probable families of models. Black outline indicates the models of interest. **B** Parameter estimates extracted from the winning family of models of young participants and second probable model of old participants. Bar height represents the grand average across old (darker colours) and young participants (lighter colours) for high predictability (green) and low predictability (orange), error bars represent *±*1 SEM.

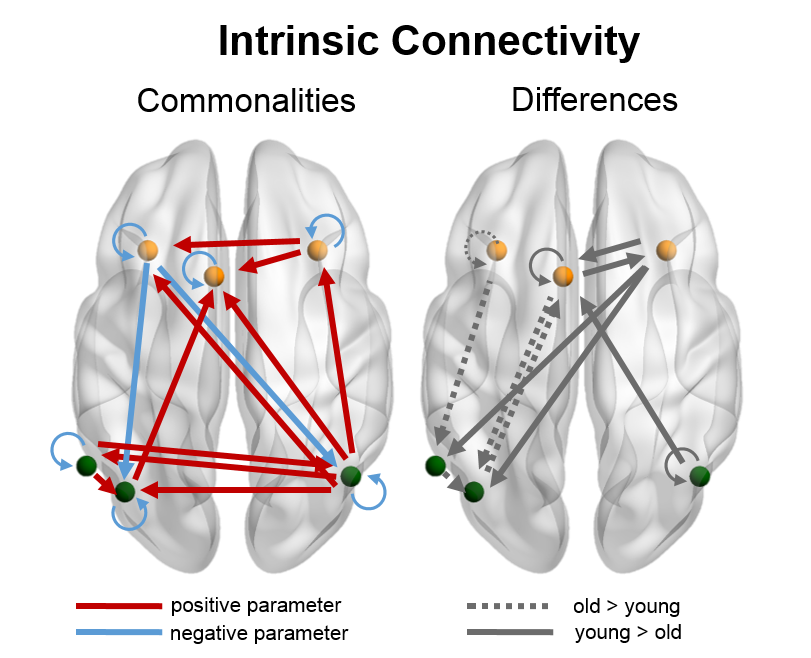

**Figure 4. DCM PEB results**. Group results of the Bayesian model average (BMA) of the intrinsic parameters (A-parameters). Left: Common intrinsic connectivity parameters across both groups, right: age-related differences in intrinsic connectivity.

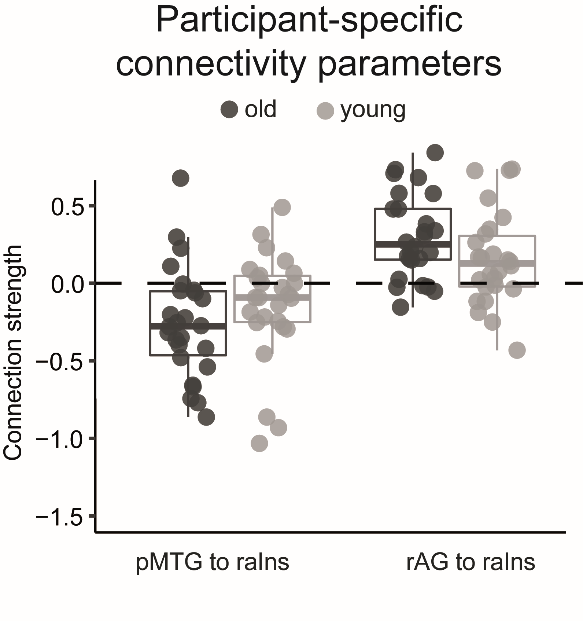

**Figure 5.** **DCM PEB results**. Individual modulatory parameters for high predictable sentences at each connection with a common effect for both age groups at a threshold of 95 % posterior probability; laIns/raIns = left/right anterior insula, pMTG = posterior middle temporal gyrus, l/rAG = left/right angular gyrus, pre-SMA = pre-supplementary motor area.

**Table 7. DCM PEB Results.** Parameter estimates from the Bayesian model average (BMA) of intrinsic connectivity.

| **Intrinsic Connectivity - Commonalities across age groups** | | |
| --- | --- | --- |
| **Connection** | **Intrinsic connectivity** | |
|  | **Parameter (Hz)** | **Pp** |
| *Self-connections* | | |
| Pre-SMA self | **-0.43** | **1** |
| laIns self | **-0.33** | **1** |
| raIns self | **-0.50** | **1** |
| lAG self | **-0.43** | **1** |
| rAG self | **-0.65** | **1** |
| pMTG self | **-0.47** | **1** |
| *Within Semantic Network* | | |
| rAG -->lAG | **0.11** | **1** |
| rAG -->pMTG | **0.06** | **1** |
| pMTG -->lAG | **0.06** | **1** |
| pMTG -->rAG | **0.10** | **1** |
| *Within Cingulo-Opercular Network* | | |
| raIns --> laIns | **0.12** | **1** |
| *Between Network Connections* | | |
| laIns -->lAG | **-0.12** | **1** |
| laIns -->rAG | **-0.13** | **1** |
| lAG -->pre-SMA | **0.05** | **1** |
| rAG -->pre-SMA | **0.05** | **1** |
| rAG -->laIns | **0.05** | **1** |
| rAG -->raIns | **0.15** | **1** |

Note: Only connections with at least one parameter exceeding a posterior probability (Pp) > .95 are shown and highlighted in bold.

**Table 8.** **DCM PEB Results.** Parameter estimates from the Bayesian model average (BMA) of intrinsic connectivity.

| **Intrinsic Connectivity - Differences between age groups (young > old)** | | |
| --- | --- | --- |
| **Connection** | **Intrinsic connectivity** | |
|  | **Parameter (Hz)** | **Pp** |
| *Self-connections* | | |
| Pre-SMA self | **0.06** | **1** |
| laIns self | **-0.13** | **1** |
| rAG self | **0.06** | **1** |
| *Within Semantic Network* | | |
| pMTG -->lAG | **-0.06** | **1** |
| *Within Cingulo-Opercular Network* | | |
| laIns -->pre-SMA | 0.02 | 0.53 |
| pre-SMA -->raIns | **0.05** | **1** |
| raIns -->pre-SMA | **0.09** | **1** |
| *Between Network Connections* | | |
| pre-SMA -->lAG | **-0.06** | **1** |
| laIns -->pMTG | **-0.08** | **1** |
| raIns -->lAG | **0.11** | **1** |
| raIns -->pMTG | **0.11** | **1** |
| lAG -->pre-SMA | **-0.09** | **1** |
| rAG -->pre-SMA | **0.09** | **1** |
| pMTG -->lAG | **-0.06** | **1** |

Note: Only connections with at least one parameter exceeding a posterior probability (Pp) > .95 are shown and highlighted in bold. For the differences between age groups, positive values indicate stronger connectivity for young adults.
